# FluoroTensor: identification and tracking of colocalised molecules and their stoichiometries in multi-colour single molecule imaging via deep learning

**DOI:** 10.1101/2023.11.21.567874

**Authors:** Max F.K. Wills, Carlos Bueno Alejo, Nikolas Hundt, Andrew J. Hudson, Ian C. Eperon

## Abstract

The identification of photobleaching steps in single molecule fluorescence imaging is a well-established procedure for analysing the stoichiometries of molecular complexes. Nonetheless, the method is challenging with protein fluorophores because of the high levels of noise, rapid bleaching and very variable signal intensities, all of which complicate methods based on statistical analyses of intensities to identify bleaching steps. It has recently been shown that deep learning by convolutional neural networks can yield an accurate analysis with a relatively short computational time. We describe here an improved use of such an approach that detects bleaching events even in the first time point of observation, and we have included this within an integrated software package incorporating fluorescence spot detection, colocalisation, tracking, FRET and photobleaching step analyses of single molecules or complexes. This package, known as FluoroTensor, is written in Python with a self-explanatory user interface.

## 1. Introduction

Single molecule methods provide a potent approach to examine macromolecular complexes. In particular, using total internal reflection fluorescence (TIRF) microscopy to visualize fluorophores, one can determine whether different specific molecules are present concurrently in a complex, and measure their stoichiometries, rates of association and dissociation, rates of conformational transitions and rates of diffusion in two dimensions^1–4^. The measurement of exact stoichiometries is a particular strength of the method, since they can be inferred by identifying successive steps in the stochastic photobleaching of individual molecules in a complex. Photobleaching has been used very widely to examine, for example, the numbers of subunits in membrane-bound protein complexes in prokaryotic and eukaryotic cells, the number of RNA or protein molecules in RNA splicing complexes and the numbers of molecules of a ligand bound to multi-subunit proteins^2,5–13^.

The photobleaching steps of dye-labelled macromolecules can often be assigned fairly easily by eye^2,14^. However, protein fluorophores have lower rates of emission, and so the signal/noise ratio is less favourable. Nonetheless, unassisted assignments can be made^11^. Assignment by eye has three disadvantages: it is subjective, and different observers may make different assignments; it is slow, which is of especial importance when the statistics require hundreds or thousands of traces to be analysed from each experiment; and it is often difficult to make assignments because observers look for plateaus on either side of a step and may not be able to judge whether, for example, there has been bleaching within the first frame or two of the recording or whether two events have taken place simultaneously or in close succession.

Several alternatives to visual inspection have been developed. One alternative is to measure the intensity of fluorescence from the single particles or spots prior to photobleaching, and divide this by the intensity of an individual (unitary) bleaching step. A number of methods have been developed to measure the unitary step, including a pairwise-difference distribution function^1^, an iterative search for statistically significant change points and a Gaussian fit to the distribution of step heights^15,16^, and the use of the last step in a bleaching curve to measure the statistical parameters of a step and then use of Bayesian methods to find models that best fit the observed curves ^17^. A related method is the use of purified fluorescent protein to provide the single step data, and comparison of these with step spacings derived by edge-preserving filters and Fourier analysis^18^ These methods are most appropriate where the step size is roughly constant, or where the numbers of fluorophores in each spot are so high that any variation in step size can be assumed to be averaged ^17^. However, the absorption of excitation light depends strongly on the orientation of the molecular dipole; dipoles aligned in the direction of propagation of the field experience a field only around 10% of that experienced by dipoles perpendicular to the surface, and the rate of emission will be correspondingly reduced^19^. Protein fluorophores on complexes that have been captured on a surface may have restricted orientations, and so the rate of emission and thus the magnitude of bleaching steps may show very wide variation among molecules and cannot be relied upon to reveal the numbers of molecules in a complex.

An alternative approach that is initially less dependent on knowledge of the unit step size is to model the bleaching time course by identifying the plateaus, or states, that flank the steps. An advantage of defining plateaus is that blinking or reactivation of fluorophores after bleaching does not affect the number of steps identified. Plateaus have been identified by hidden Markov models^20^ or by measuring the mean and standard deviation for short segments of the curve, followed by iterative steps in which the segment was expanded^21^. Recursive binary segmentation has also been applied, with Student’s t or other statistical tests being used to maximise the difference in the means between the halves. In one recent method, this was followed by k-means clustering of segments, and then the use of a Viterbi algorithm to determine the most probable sequence of states^22^. These methods still required the amplitudes of the steps to be relatively consistent, and they perform better with increasing numbers of data points in each plateau. We have used a related method involving a Bayesian step point detector, requiring only a minimum but still arbitrary step height^5,9,10^. A serious limitation with all these methods is that bleaching is stochastic, i.e., the probability is the same for every photon absorbed. Thus, more molecules will bleach in the first time-frame than in any other, but these will be missed in any method based on detecting statistical plateaus. Moreover, it is possible that all the methods summarised above will tend to lose molecules that bleach very rapidly or that have very small step heights, implying that there might be inadvertent selection of molecules with a restricted range of dipole orientations.

Deep learning could bypass some of the limitations or assumptions required in most of the above methods, if the training set were to include stochastic bleaching events and appropriately distributed step sizes. A program incorporating convolutional neural networks has been reported that performed well on samples labelled with dye or protein fluorophores, and reduced the time required for processing by two orders of magnitude^23^. However, in this case each state of the dataset used for training and testing included at least five time points, which would compromise its ability to detect bleaching in the first frame or very close frames. We describe a similar program here that enables these events to be detected accurately. Moreover, bleaching steps can be detected even with a signal/noise ratio as low as 1.1, when an accuracy of 75% was achieved. We have incorporated this programme into an integrated package for the detection of spots, colocalization, step measurement and downstream analysis, FRET and 2D tracking. We anticipate that this package will be useful in a wide range of single molecule experiments.

## 2. Methods

### 2.1 Image Processing

The images acquired from the microscope were initially enhanced by summing the first 20% of frames for each channel to improve the signal-to-noise ratio of fluorescent foci. Additionally, the maximum projection was taken also to avoid missing foci from fluorophores that bleach rapidly. These projections were combined together and enhanced. Two alternative enhancement techniques were used. One is described in Supplementary Figures 1 and 2 (more detail in Supplementary Methods); the other is a wavelet transform. The correlation of the wavelet and image was computed for every pixel such that the output image is zero for all regions apart from fluorescent foci where the overlap integral was high.

### 2.2 Detection of Fluorescent Foci

Fluorescent foci (spots) were detected in a four-step algorithm involving a number of thresholds; the steps are shown in Supplementary Figure 3. The enhanced image was split into a grid of 8x8 pixel boxes. Each potential spot was compared with a precomputed kernel to discard aberrant bright regions such as noise or dead pixels for computational efficiency. The remaining foci are then fitted with Gaussians and kept or rejected based on fitting criteria such as width, eccentricity, and residual.

### 2.3 Correcting for Chromatic Aberration

To maintain focus when switching from 640 nm excitation to 488 nm excitation, the stage controller compensates in the z axis. This very minor change in z height between the channels, in tandem with lateral and spherical aberration, results in a slight difference in perceived magnification between the channels as shown in Supplementary Figure 4. A linear transform to correct for chromatic aberration is defined by equations 1a and 1b below^9^:

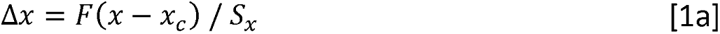

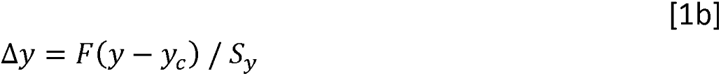

where Δ*x*,Δ*y* is the shift in coordinates of a spot due to chromatic aberration, *F* is the shift factor which scales the whole transformation based on the wavelength difference (*F*=1 for 640 nm ➔ 488 nm and 0.35 for 640 nm ➔ 561 nm corrections), (*x, y*) are the coordinates of the spot, (*x_c_, y_c_*) are the coordinates of the origin of the transformation (where spots perfectly superimpose), and (*s_x_ , s_y_* ) are scale factors of the stretch and are equal to the distance from the origin of the transformation which results in a 1 pixel shift in the respective axis. Note that here we refer to the channels by their excitation wavelengths, whereas the chromatic shift correction factors were calculated from peak emission wavelengths.

### 2.4 Calculating Fluorescence Intensity Time Traces

Fluorescence intensity time traces were calculated from the raw data without enhancement. The background was subtracted locally around the spot for each frame as shown in Supplementary Figure 5. The background subtraction ensures that if all the fluorophores in a fluorescent focus bleach, the mean intensity of the trace after that point will be zero. This is important as the neural network will only confidently assign stoichiometries where all the fluorophores have bleached. If one or more fluorophores don’t bleach within the recording time, the intensity will be above zero and the stoichiometry will be undecided.

### 2.5 Neural Network Architecture

The architecture of the convolutional recurrent neural network (CRNN) is shown in figure 1. The fluorescence intensity trace was convolved with a set of filters learned during training. This produces a set of feature maps which were then further convolved deeper in the network by even larger sets of filters. The feature maps were then passed to a long-short-term-memory (LSTM) layer. This increases prediction accuracy as the LSTM can ignore photo-blinking by the presence of features such as an upwards step when the fluorophore switches back on followed or preceded by a photobleaching event of a similar magnitude^23^. The output of the LSTM was passed to a multilayer perceptron which classifies the trace as having zero, one, two, three, four, five or more, or an undefined number of steps based on the combinations of features found by the LSTM.

**Figure 1.**
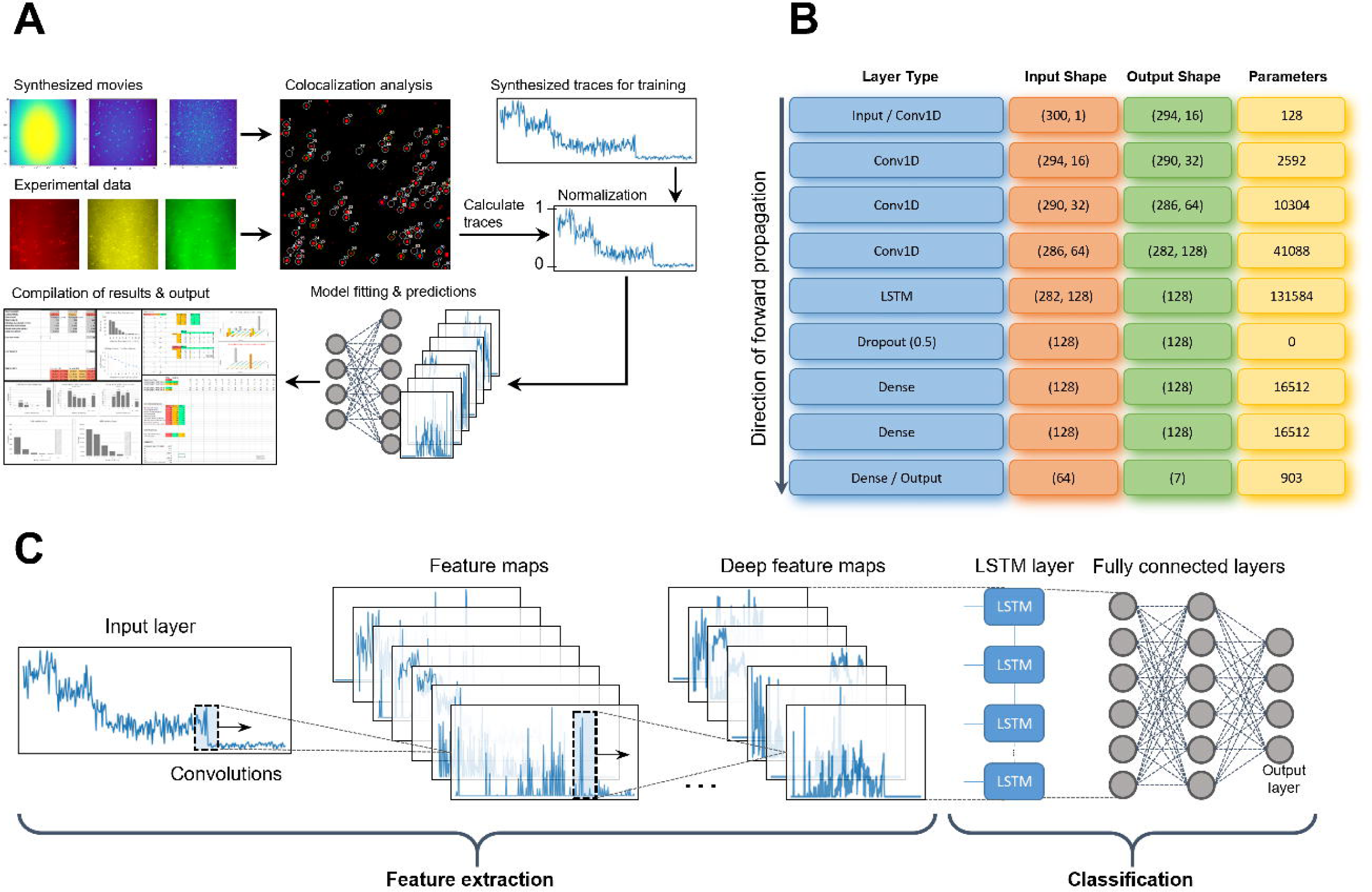
(A) Scheme showing data processing pipeline of the FluoroTensor package. Synthesized traces are used to train the CRNN model based on the properties of fluorophores in use. The full pipeline is then tested with synthesized single molecule movies. A projection of each channel is enhanced, fluorescent foci detected, colocalization between channels is mapped, and time intensity traces are calculated based on methods covered later in this section. Traces are then passed to the CRNN which predicts the number of photobleaching steps, and a moving average fitting tool takes this as an input and attempts to find the most likely time points where photobleaching occurs. The results are then compiled into an excel spreadsheet. **(B)** Table of layers in CRNN along with the shapes of their respective inputs, outputs, and number of trainable parameters. **(C)** Schematic diagram of the layers of CRNN.

A dropout layer was added between the LSTM and the dense layers to reduce overfitting ^24,25^. During training, 50% of neurons in the dropout layer were ignored at random during forward propagation. This prevents the network from over-relying on specific weights which effectively regularizes the model.

### 2.2 Network Training

CRNN models were implemented in TensorFlow 2.7 using the Keras API. Models were trained on a Gigabyte GeForce 1070 and took approximately 24 hours to converge. To prevent overfitting, real-time validation was computed on an unseen validation set during training. Model weights were only saved if the validation sparse categorical accuracy had improved from the previous best epoch. The model was optimised using Adam with a learning rate of 0.0001 and the sparse categorical cross entropy loss function.

The CRNN architecture was trained independently on three separate artificial datasets to create three models tailored to different fluorophores, namely Cyanine 5 (Cy5), monomeric Cherry (mCherry), and monomeric enhanced green fluorescent protein (mEGFP). Cy5-like traces were simulated with constant step heights ± 20% allowing for some variation, and mCherry / mEGFP – like traces were synthesized with intensities drawn from a cos^2^ distribution^26^. Each training set was also synthesized over a range of signal-to-noise ratios to account for almost every case typically encountered with our experimental setup. Supplementary Figure 6 shows examples of synthesized Cy5 traces and signal-to-noise ratio distributions of the training sets for each type of fluorophore. Traces were synthesized by first generating an idealized trace with stochastic photobleaching where the probabilities of virtual fluorophores based on the number of photons absorbed. Noise was then superimposed on the ideal trace. The simulation method is covered in more detail in Supplementary Materials. Average signal to noise ratios (aSNRs) of the steps of a trace were calculated using equation 2^23^:

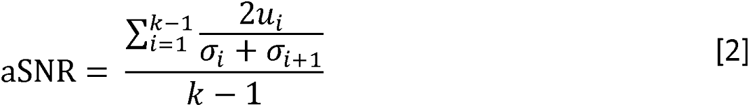

where *k* is the number of plateaus (number of steps + 1), *u_i_* is the absolute difference between the mean of the i^th^ plateau and the (i+1)^th^ plateau, and *σ_i_* is the standard deviation of the i^th^ plateau.

### 2.7 SM Tracking

Initially each frame was enhanced using a high pass filter. This was achieved by convolving the frame with a normalized Gaussian kernel (128x128, σ = 32, with padding) which acts as a low pass filter which was then subtracted from the original (see Supplementary Figure 7 for example). The signal-to-noise ratio of the spot in the raw image data was calculated as the background subtracted mean intensity of the spot masked at the full-width-half-maximum of its Gaussian fit divided by the standard deviation of the surrounding background noise (see Supplementary Figure 8).

Molecules are detected on a framewise basis using the same detection algorithm as previously described for colocalization experiments. The fitting criteria are more relaxed for tracking experiments owing to the lower signal to noise ratio of individual frames compared to averaged frames in colocalization experiments. Tracks are connected by a temporal nearest neighbour algorithm. A spot was deemed to be the same molecule from a previous frame if it was the nearest to the previous position and within a threshold distance to prevent tracks connecting to other molecules across the image. If a molecule photo-blinks and is not detected for several frames, the algorithm will continue tracking it when it comes back within a small (typically 1 frame) time window and if it was still within the threshold distance, but otherwise it was treated as the beginning of a new track.

The diffusion coefficient for each track was calculated by taking the linear regression of the MSD plot and dividing the gradient by 4 for 2D diffusion. The method for calculating the MSD for different lag times (tau) is shown in equations 3a/b^27^.

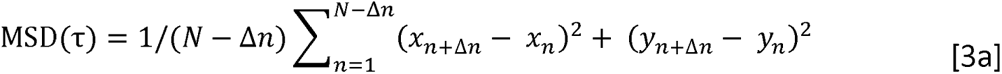

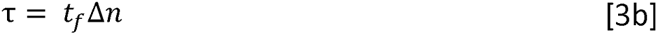

where MSD(τ) is the mean square displacement at lag time *τ*, Δ*n* is a nonzero positive integer equal to the number of frames between the position of the spot at frame *n* and its position at frame *n*+Δ*n* , N is the total number of frames, *x* and *y* are the coordinates of the spot in the respective frame, *t_f_* is the frame duration. The MSD was then plotted against *τ* for every possible Δ*n*.

### 2.8 Sample Prep

Proteins fused to mCherry and mEGFP were expressed in HEK293T cells. Nuclear extracts were prepared and samples were injected into chambers on the cover slip as describe^5,10^. Images were acquired as described ^9^ with a minor change – the number of frames recorded was standardized to 300 in line with the neural network’s input domain. For the analysis of 2D diffusion, dye-conjugated oligonucleotides were conjugated to a hydrophobic moiety and injected onto a hydrophobic surface as described (Santana Vega et al, manuscript in preparation). Images were acquired by TIRF.

## 3. Results

The overall purposes of this work were to use deep learning to develop an improved method for detecting the numbers of bleaching steps in time courses of fluorescence and to embed it in an integrated software package that could facilitate and accelerate much of the data analysis associated with TIRF microscopy. Prior to the extraction of time course data, images need to be processed to reveal faint signals and, in the case of multicolour fluorescence experiments in which molecules or complexes might contain more than one fluorophore, colocalized signals need to be identified. These steps are followed by extraction of the intensity data for each spot detected and identification of the number of steps in which each colour bleached (Figure 1).

### 3.1 Detection of fluorescent molecules or complexes

Since protein fluorescence intensities can vary widely, FluoroTensor incorporates a set of operations designed to detect signals over a wide range. The detection of dim signals is important to avoid under-representation of weak emitters, which are likely to contain only a single fluorophore. Initial trials showed that a high-pass filter did not produce an image with a satisfactory background removal and sufficient contrast for weak emitters such as mCherry (Figure 2). Instead, the signal/noise ratio is improved by summing the frames for a user-specified proportion of the acquisition (usually the first 20%, which reduces the inclusion of frames acquired after bleaching of the spots). In addition, the maximum pixel values in this region are taken, to improve the visibility of spots that bleach rapidly. The summation and maximum values are combined and normalised. This image is enhanced in either of two ways (as described in methods).

**Figure 2.**
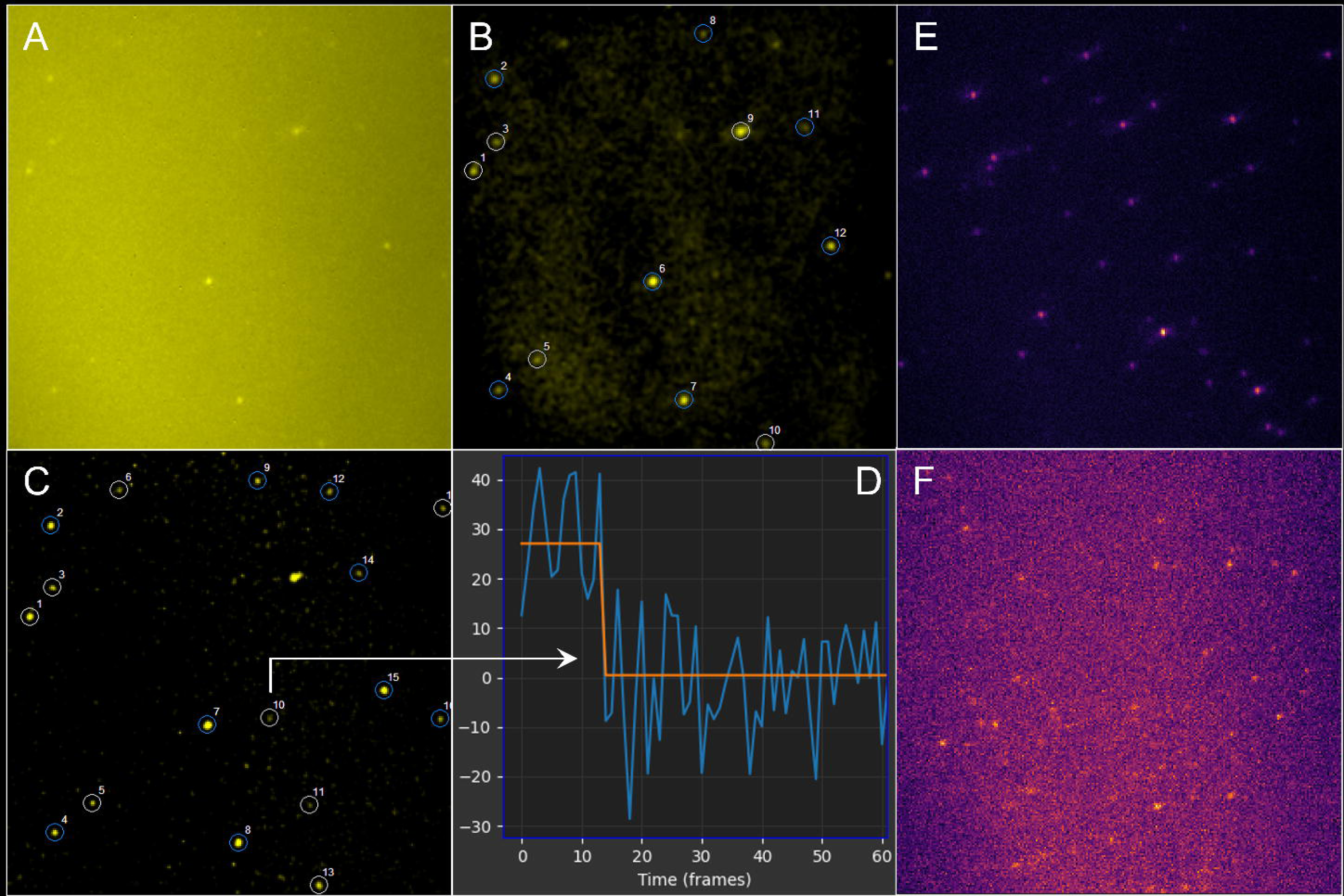
(A) A composite image obtained by taking the mean of the first 20% of all frames in the mCherry Channel and combining it with the maximum projection of those same frames. **(B)** The composite image enhanced using a high pass filter and showing the spots that were detected. **(C)** The composite image enhanced using the method detailed in section 2.1 (figure 1) and the spots detected. Note how spot 9 kept in the image enhanced by the high-pass filter is rejected here owing to the higher contrast of our enhancement resolving a second weaker spot overlapping it. **(D)** A zoom of the start of the trace of spot 10 proving that the spot detected was a real single mCherry molecule and was not an artefact of the enhancement. **(E)** A single frame from a movie of Cy5-labelled RNA molecules. **(F)** A single frame of mCherry-labelled U1A proteins from the same experiment as (E), note the much lower signal to noise ratio and thus the need for a powerful enhancement technique. Comparing (A) to (F) we can see that a single time bin (100ms) of Cy5 dyes has a much greater signal to noise ratio than a summed stack of 60 100ms time bins of mCherry fluorescent proteins. (Note that while (A) and (E) have different colour maps they are both still perceptually uniform.)

Spots are detected using a pre-computed Gaussian kernel and calculation of the residual between the spot and the kernel. Those spots that satisfy this criterion are fitted by a Gaussian curve to calculate the coordinates of the spot with sub-pixel precision. This procedure involves less computation than the alternative of trying to fit all possible regions of the field of view to a Gaussian. Any remaining false positives are discarded during downstream analysis of the fluorescence trace due to the absence of photo-bleaching steps.

The colocalization of spots in images acquired at different wavelengths is often complicated by chromatic aberration. This arises from the different angles of refraction at those wavelengths. resulting in a difference in focal depth between the channels. This effect is generally counteracted either by the use of fiducial beads in the sample or by prior calibration of the system (equations 1a and 1b) with beads or dual-labelled molecules. However, we observed that, when the stage was automatically moved to a new area on the slide, in some cases the autofocus would keep the field of view in focus but at a slightly different focal height, resulting in a change in chromatic aberration with slightly different parameters. This change could result in some spots not being identified as colocalized. To solve this, we incorporated an additional optional solution in which an optimizer is implemented directly into the program to solve the parameters to equations 1a/b for each field of view during automated analysis. The approach is to solve the parameters for maximum colocalization. However, the colocalization count is not a continuous function of the parameters and as such, unsuitable for optimization by commonly used solvers such as Simplex. The first step is to test an array of transformation origins and scale factors to find an approximate set of parameters which maximises the number of colocalized spots via grid search. Next, a Simplex solver minimizes the distances between colocalized spots, taking the crude parameters as an input and refining them. A minimum of 4 colocalized spots is required for this to work reliably and spots should be sparsely populated, especially if the colocalization percentage is low, to discourage the solver from attempting to colocalize randomly proximal spots.

### 3.2 Determination of stoichiometry by a convolutional neural network

The architecture was designed to be efficient as well as accurate. The FluoroTensor platform enables users to quickly train their own models from this architecture. The architecture is small, totalling only 220K parameters, and yet it shows improved accuracy (Figure 3) when compared with larger models ^23^. Our models were trained independently on 3 different datasets totalling 18 million synthetic traces. Each model took 24-48 hours to train on an NVIDIA GTX 1070 GPU using a modified Keras generator to load datasets of 240,000 traces sequentially during training. The use of synthetic traces was mandated by the need to know the number of steps in each trace and to ensure equal representation of traces with the different numbers of steps being classified. A plateau length restriction was placed on traces with very low signal to noise ratios (Supplementary Figure 9), such that if two bleaching steps were indistinguishable from a single step, the intermediate plateau was extended, and no other fluorophores were allowed to bleach in the simulation until the intermediate plateau becomes resolvable. Statistically only a small proportion of the training data has these extended plateaus based on the SNR distribution. Of great importance to note is that these restrictions were only imposed on the training dataset to prevent the model from overpredicting steps. All synthetic testing data was simulated with stochastic bleaching, allowing for simultaneous bleaching events that would be present in real data unless explicitly stated otherwise.

**Figure 3.**
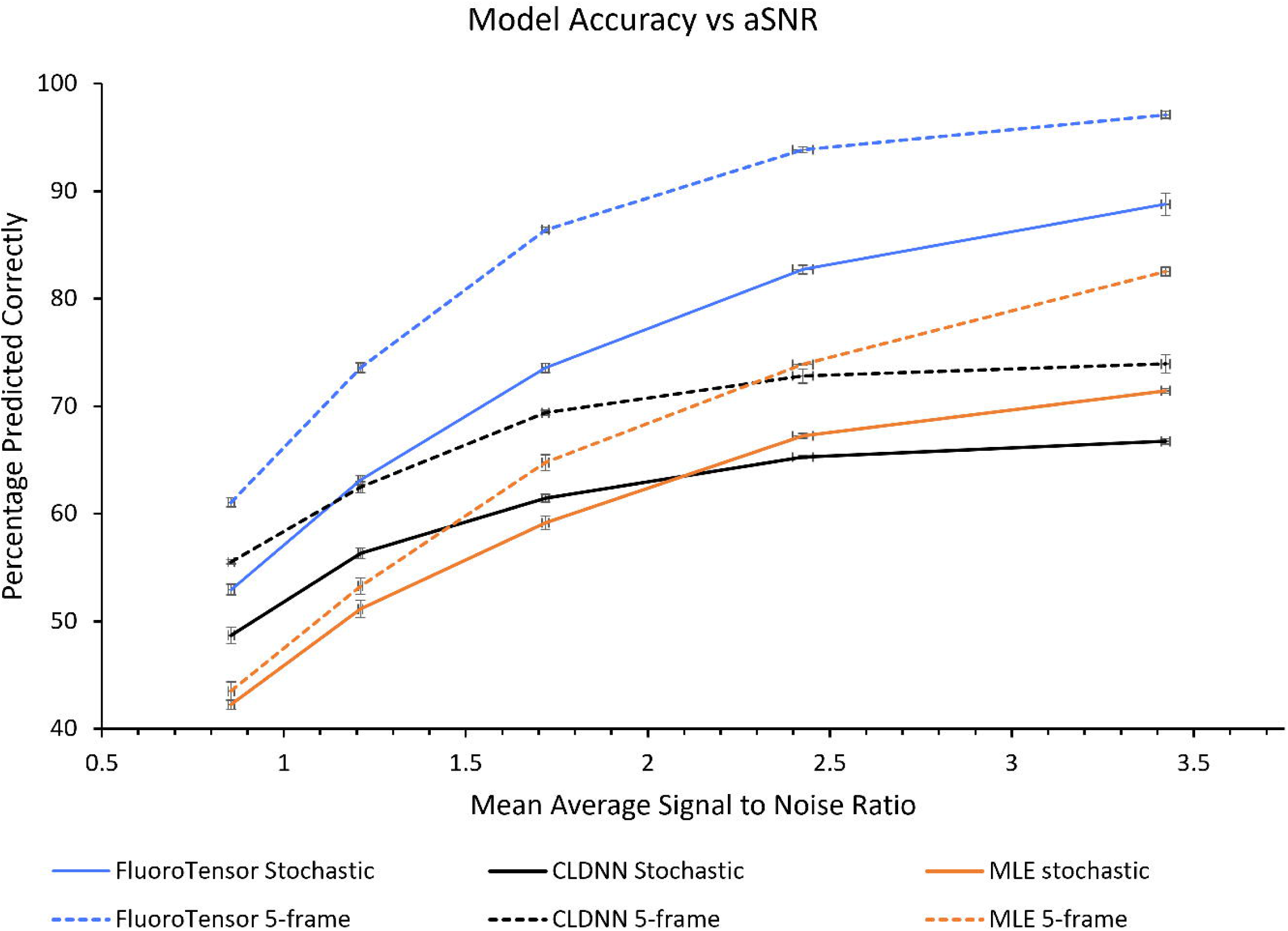
A comparison of the FluoroTensor models with CLDNN and MLE for stochastic bleaching traces and traces with a minimum of 5 frames per plateau as described in Xu et al 2019 for CLDNN. Comparisons were made between prediction accuracy of each model for datasets with a range of signal to noise ratios (more detail in Supplementary Figure 14).

The network was tested using simulated data for organic fluorophores that lacked constraints and therefore included some simultaneous bleaching events. These tests were done on sets of data with a range of signal/noise ratios and various mean fluorescence intensities (Supplementary Figure 10). The accuracies recorded increased with the signal/noise ratio, as expected, from 74% at a ratio of 1.105 to 98.1% at a ratio of 8.853. Accuracy will never reach 100% due to the inherent uncertainty of simultaneous photo-bleaching events regardless of the signal to noise ratio. To gain insights into the feature extractor part of the neural network, the 128-dimensional feature vectors (the outputs of the LSTM layer of the model) of a dataset of 20,000 traces were processed by t-distributed stochastic neighbour embedding (t-SNE) dimensionality reduction algorithm to produce a 2-dimensional map clustering the feature vectors by similarity (Supplementary Figure 11). Comparing the feature vector t-SNE plots with the t-SNE plots built from the traces themselves (Supplementary Figure 12) it is clear that the convolutional and LSTM layers of the network are working as intended, reducing the extremely high intra-class variance of the traces significantly into highly conserved feature vectors which can then be decoded by the multilayer-perceptron classifier. Crucially, the intra-class variance is hugely reduced while the inter-class variance is increased, i.e., feature vectors are highly correlated between members of the same class and highly uncorrelated between members of different classes.

The experimental setup and operating procedure described here have been optimised for recordings of 300 frames (time points in the traces) for each colour channel. Convolutional Neural networks have a fixed size input domain and thus any trace entering the model needs to be resampled to 300 frames to match the 300 neurons of the input layer. As shown in Supplementary Figure 13C, resampling a trace from 100 frames to 300 frames using a linear interpolator introduces extra time points between plateaus. Once the number of interpolated points is on the order of the number of frames of the shortest plateaus, the neural network is unable to distinguish the interpolated frames from a true plateau and begins to overpredict the number of steps in the trace resulting in significantly reduced accuracy, as shown in Supplementary Figures 13A and 13B. Interestingly, when the bleaching half-life is sufficiently short such that a number of steps would be missed due to simultaneous bleaching events, the accuracy of the 200-frame dataset interpolated to 300 frames was slightly greater than the accuracy of the native 300-frame dataset. This effect can be attributed to overprediction caused by resampling the 200-frame that was compensating for the underprediction due to simultaneous bleaching events. The inverse linear correlation between the probability of a fluorophore bleaching per frame and the accuracy of the neural network is shown in Supplementary Figure 13D. Almost all errors were due to underpredictions where the network is unable to distinguish bleaching steps with more than one bleaching event. For this reason, it would be advisable to choose a frame rate and laser power that maximises the mean time interval between bleaching events while also ensuring that bleaching is likely to be complete for all likely numbers of bleaching steps by the end of the 300 frames.

### 3.3 Comparisons of performance

To compare the performance of FluoroTensor with a previous neural network model (CLDNN^23^) and with a statistical method^9,28^, time courses were simulated for complexes containing 0 to 4 protein-like fluorophores across a range of mean signal/noise ratios (mean bleaching half-life was 40.5 frames, see Supplementary Figures 14 and 15). The percentage of complexes predicted correctly by each of these methods is shown in Figure 3. In addition to simulations of stochastic bleaching, a second set of simulations was made with a minimum of 5 frames between each bleaching step. The results show that FluoroTensor was significantly more accurate for each dataset than the other two methods. Interestingly, in both data sets the CLDNN model performed better than the statistical method, based on maximum likelihood estimation^29^, at low signal/noise ratios whereas the MLE method was superior at higher signal/noise ratios.

A further comparison was done to analyse whether the abilities of the methods to determine the distributions of complexes with various numbers of fluorophores would indicate the reasons behind the differences in performance. Three SM TIFF stacks were simulated with virtual complexes formed with a number of dye-like fluorophores drawn from binomial distributions based on the numbers of molecules in a complex (2, 3 and 4) and the proportion of these that were labelled (p = 0.5, 0.4 and 0.6 respectively; Figure 4A). Complexes were modelled as Gaussian functions with sigma = 1.3 and an amplitude equal to the sum of the intensity of the virtual emitters. The fluorophores were allowed to bleach stochastically. Gaussian noise was added to simulate dark current, a non-uniform autofluorescence was added to each frame and shot noise was modelled with a gamma distribution to account for EMCCD gain.

**Figure 4.**
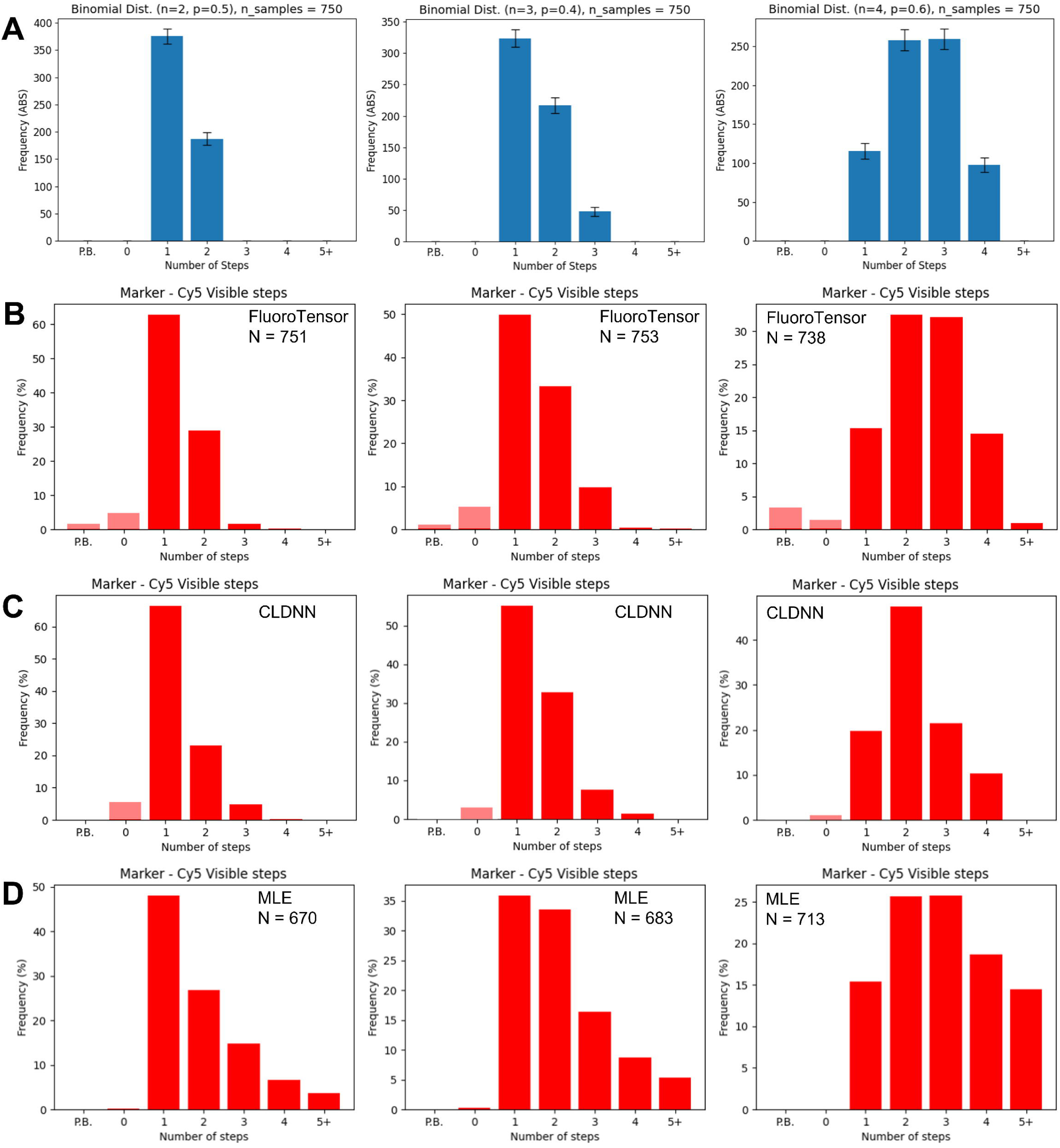
(A) Three different binomial distributions. Consider the number of trials, N, as being the number of bound proteins in a complex and the probability of success, p, the ratio of exogenous FP tagged protein to endogenous untagged protein. The binomial distribution represents the distribution of photobleaching steps of fluorescent foci of complexes with these stoichiometries and labelling proportions. **(B)** Simulated data with these distributions was analysed automatically by FluoroTensor including locating the foci in the SM movies, calculating traces and using the neural network models to predict the number of photobleaching steps. **(C)** The same analysis on the same traces as (B) but using the CLDNN model to predict steps. **(D)** The distributions of steps found by a maximum likelihood estimator in our previous SM analysis software, Auswerter, developed in MATLAB (Jobbins et al., 2022).

The three stacks were analysed automatically by FluoroTensor to identify the spots and the stoichiometries of the simulated complexes. The distributions observed matched the expected binomial distributions (Figure 4B). These distributions were then used to predict the numbers of molecules in the complexes, producing answers of 2 (with a confidence of 78%), 3 (with a confidence of 83%), and 4 (with a confidence of 80%). The analysis was repeated on the same set of extracted traces using the computation neural network^23^, with the results shown in Figure 4C. This underestimated the complexes with three or four steps, and the number two was overestimated, possibly because close bleaching steps were not assigned to distinct molecules. The TIFF stacks were also analysed using a MATLAB suite of programs that enabled spot detection and assigned stoichiometries using an automated step detection algorithm based on maximum likelihood estimation (MLE; Figure 4D)^9,28^. In contrast to CLDNN, this analysis tended to overestimate the abundance of complexes with higher stoichiometries, possibly because of the signal/noise ratio (distribution shown in Supplementary Figure 16).

Apart from the accuracy, there are two further advantages of FluoroTensor compared with statistical methods: the speed of the analysis and the ability to correct automatically for chromatic aberration. An example illustrating these advantages is shown in Supplementary Figure 17A. A dataset from TIRF microscopy of a sample of nuclear extract containing regulatory splicing complexes with fluorescently labelled components of interest and fluorescently labelled RNA was analysed both with FluoroTensor (Supplementary Figure 17A) and the MLE-based method (Supplementary Figure 17B). The FluoroTensor automated run was complete within 45 minutes of elapsed time on the dataset. No calibration file was used for chromatic aberration. Instead, the built-in optimizer detailed in section 3.1 solved the correction parameters for chromatic aberration and built up the dataset of colocalized traces. This process was entirely unsupervised apart from the final step where the program was prompted to detect the steps using the neural networks and then export the data to a preformatted excel template. In the MATLAB program, the analysis needed to be supervised with manual corrections to the spots detected and the process took over 16 hours. The software also relied on a calibration file which had pre-solved parameters from a set of calibration movies meaning that it does not accommodate any variations in the chromatic aberration from one file to the next due to subtle changes in focus. Errors in steps detected by maximum likelihood compared to the neural network model are shown for some traces in Supplementary Figure 18. These errors are thought to be made because MLE requires a threshold step size. Since these step sizes are highly variable in FP-like traces some weaker steps will be missed and brighter ones overcounted due to the necessity of setting a low enough threshold as not to miss the weaker emitters. Thus, we propose MLE could be a viable alternative for organic dyes with constant step sizes.

### 3.4 Localization Accuracy of 2D Tracking Algorithm

The spots representing macromolecular complexes are not always stationary. TIRF microscopy is used to track components undergoing lateral diffusion in membranes^30–32^ or other surfaces onto which complexes are adsorbed without covalent tethering. In such cases, estimates can be made of the diffusion coefficients of the complexes, and they may reveal heterogeneity in the surface or the interactions of the fluorescent component. However, the accuracy with which the position of a spot can be determined in a single frame is critical. In FluoroTensor, the background in each frame is removed with a high pass filter, and the location of each spot is determined as described above, beginning with a pre-computed Gaussian kernel. The analysis of simulated spots (see Materials and Methods) showed, as expected, that the mislocalization error shows a strong dependence on the signal to noise ratio of the spot in each frame (Supplementary Figure 19, 20, and 21). With a mean SNR of less than ∼2.9, some spots almost disappeared into the noise in some frames, leading to outliers with very high mislocalization. Measured spots from real single molecule data, collected from Alexa-647 ® tagged oligonucleotides diffusing on a surface typically had a mean SNR of 5 or higher, which according to our findings would have a mean mislocalization of ∼20nm on our SM TIRF microscope, with 99.7% of measured positions being localized within 50nm within a time bin of 100ms. Tracking is done by connecting each spot in a frame with the spot in the closest location in the succeeding frame (Figure 5). A maximum jump distance per frame is set based on observed motion of the moving particles to ensure valid connections of a particle one frame to the next.

**Figure 5.**
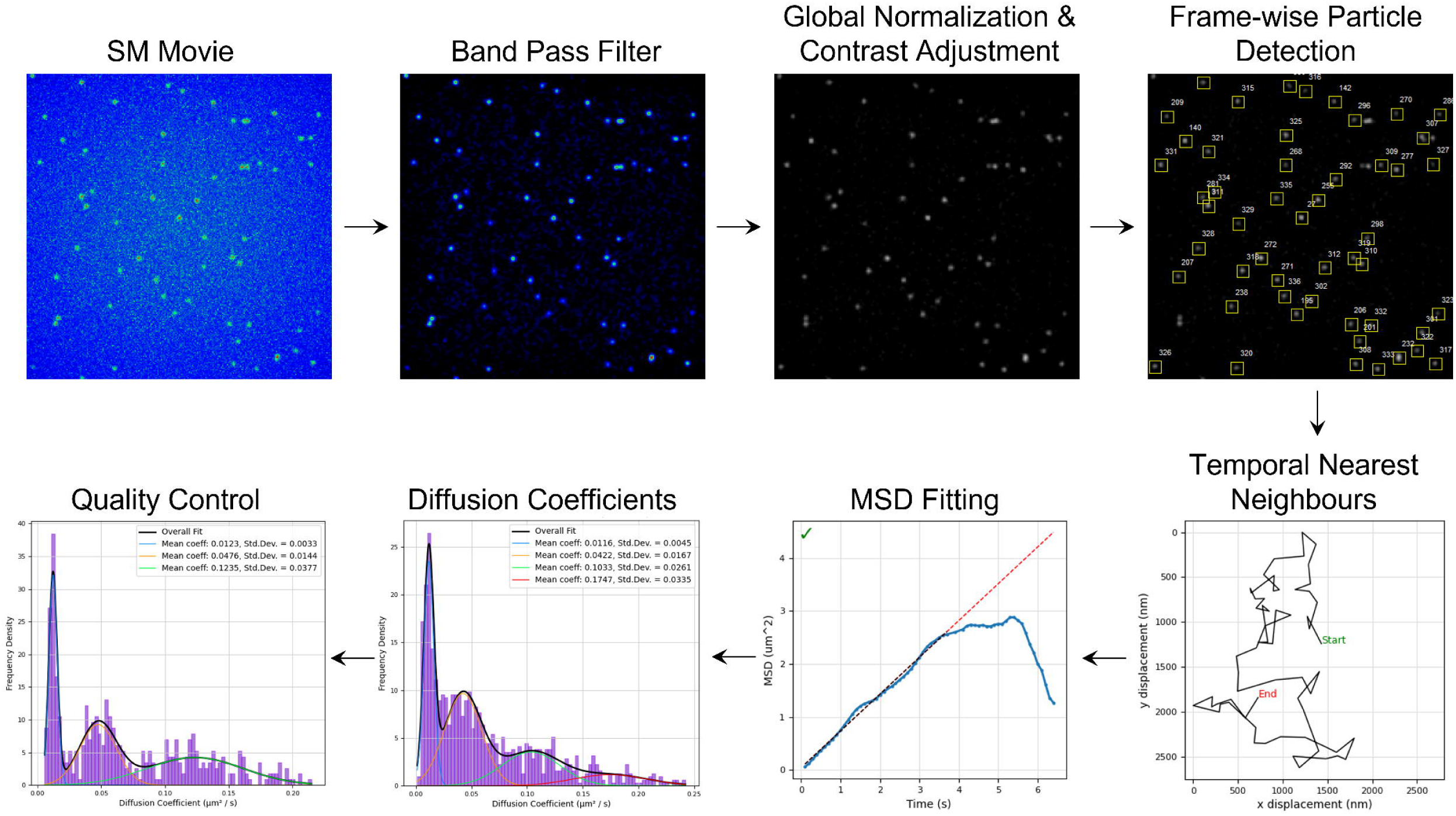
The data analysis pipeline for single molecule tracking in FluoroTensor. First, images are enhanced via high-pass filtering. Then, the movie is normalized to the brightest pixel throughout to a range of 0-255. Molecules are detected frame by frame using the same algorithm as for colocalization analysis (see Supplementary Figure 3). Spots are connected to their nearest neighbours in consecutive frames with constraints to ensure tracks are topologically linear as opposed to branching. MSDs are calculated track by track (see Supplementary Figure 22) and diffusion coefficients are generated and their distributions plotted and fitted with a Gaussian mixture model. Quality control is performed by rejecting tracks with high variance in the MSD fit which improves resolution of distributions of systems of diffusing molecules with multiple diffusion rates.

### 3.5 Estimation of Error in MSD Fit and Diffusion Histograms

Mean square displacement is calculated according to equation 3a for each track found by the program. To ensure accurate distributions of diffusion coefficients of molecules in the sample to a degree where heterogeneity in motion can be distinguished, the quality of the MSD fit must be assessed over its range and maximised. As Δn→N (equation 3a), the mean displacement no longer represents the RMS Brownian diffusion distance of a model random walk of indefinite length. This results in severe deviation of MSD vs τ from a linear fit, especially when the end-to-end distance of the track is far from the expected RMS separation (Figure 5; Supplementary Figure 22B). For this reason, only the initial portion of the beginning of the MSD plot is fitted. The proportion of the MSD plot to fit is decided on a track-by-track basis by calculating a linear regression for proportions of the data points in the MSD plot. The fitted region of the plot ends where the derivative of the coefficient of determination (R^2^) becomes negative. The final MSD fit is taken as the linear regression of all points of the MSD plot before this point. The standard deviation of the diffusion coefficients for all fitting percentages up until the point at which the derivative of R^2^ becomes negative is calculated to be used as a rough estimate of the error in the diffusion coefficient calculated by the final MSD fit (Supplementary Figure 22B/C).

To resolve the diffusion coefficients of a heterogeneous mixture, all tracks where the standard deviation as a percentage of the diffusion coefficient was greater than a specified value are rejected. This strategy results in histograms where the distribution of diffusion coefficients in a heterogeneous mixture of diffusing molecules with two or more distinct diffusivities is more easily resolved in a Gaussian mixture model than the unfiltered distribution. Figure 6 shows the effect of filtering by rejecting tracks with various estimated fitting errors. The most accurate fit was obtained with the most stringently selected data, albeit at the expense of a reduced number of data points (Figure 6; Supplementary Figure 22, D-F). This technique will be of particular use when tracks are short due to transient binding to a surface or short bleaching times of the fluorophores.

**Figure 6.**
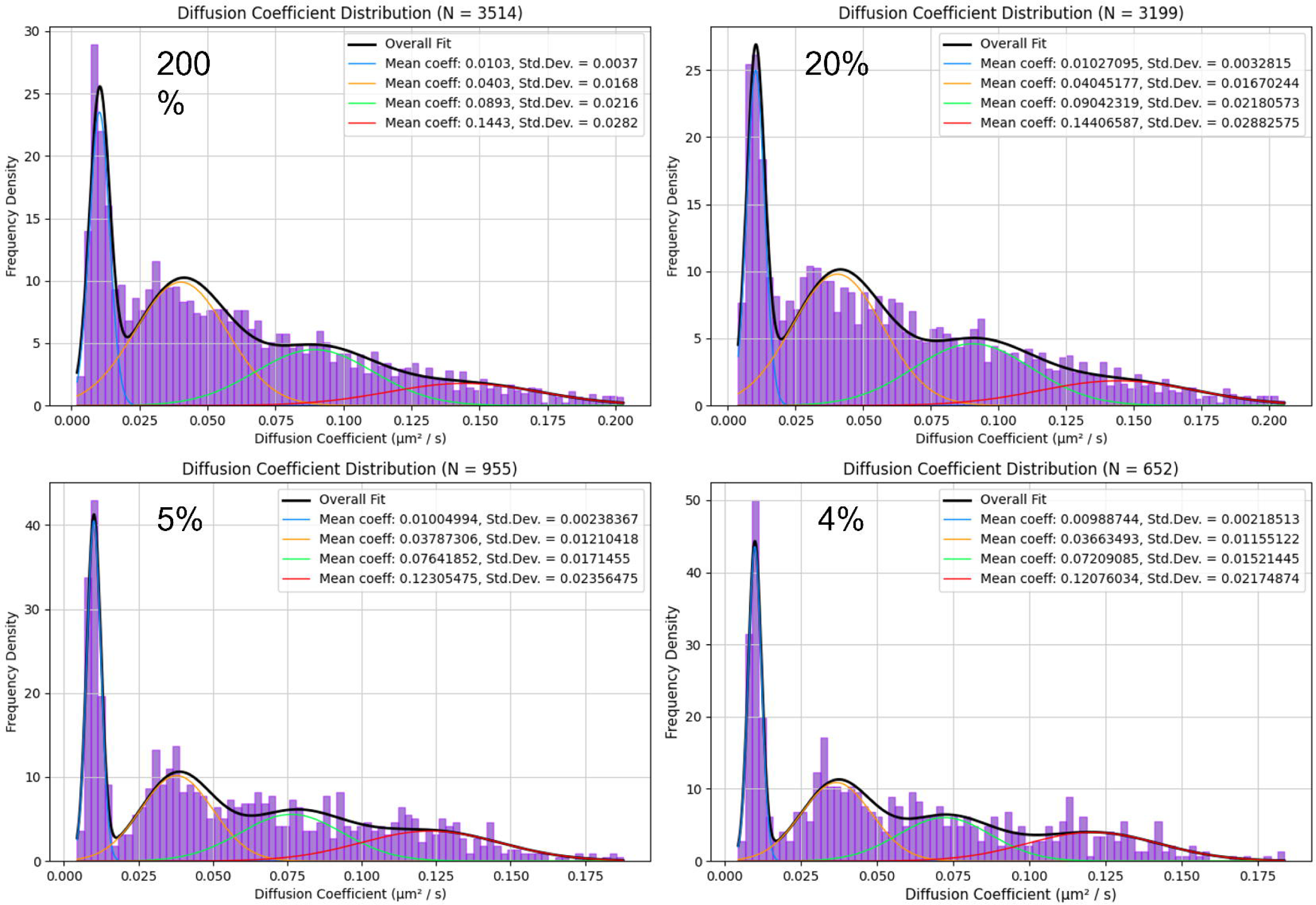
A distribution of diffusion coefficients for a system with molecules diffusing at 4 distinct diffusivities. The standard deviation of the MSD plot gradients during fitting is expressed as a percentage of the diffusion coefficient of each track. The track is rejected from the distribution of diffusion coefficients if the percentage standard deviation is greater than the respective thresholds shown (200%, 20%, 5%, and 4% respectively. Setting the threshold lower rejects a greater number of tracks. Rejecting tracks with higher variance in the MSD fit deconvolves the distribution and allows for accurate fitting of the components via Gaussian mixture model. The ground truth diffusivities simulated were 0.010, 0.035, 0.070 and 0.120 µm^2^ / s.

#### Fluorescence Resonance Energy Transfer

FRET involving two components is detected using the same Gaussian mixture model that was used above for resolving the diffusion coefficients in a heterogeneous mixture. This detects FRET states lasting for four or more frames, and, assuming a first order process, produces rate constants for the two steps. An example is shown in Supplementary Figure 23.

## 4. Discussion

We have presented a versatile new software package written in python for single molecule total internal reflection fluorescence (SM TIRF) microscopy. The FluoroTensor software package will be of particular use in two main areas. Primarily this software was designed to provide an automated analysis pipeline for single molecule multicolour colocalization microscopy with automatic correction for chromatic aberration, high contrast enhancement capable of resolving extremely weak signals from fluorescent proteins and the ability to detect photobleaching events in consecutive frames, i.e., without the requirement for identifying a statistical plateau. Within FluoroTensor, we present a neural network architecture for determining the number of photobleaching steps in the fluorescence traces extracted in the aforementioned automated analysis of SM data with 96% fewer parameters than the previous state of the art model for this purpose^23^ and with improved accuracy. The model is much more efficient to train, and our pre-trained versions are not limited to a minimum number of frames per plateau. A primary aim for this project was creating a ‘plug and play’ platform for these neural networks such that end users could train their own models based on our architecture on custom datasets, or even develop the architecture further with a relatively simple python script. Alternatively, users may finetune one of our pretrained models with their data. In addition, The SM tracking extension is well suited for tracking SM diffusion in noisy conditions where particles of interest are transient or photo-bleach rapidly. The tracking add-in which can be launched from within the main GUI also allows for automated chaining of raw data which is analysed sequentially and saved as a document file that can be reloaded and exported to Excel. We believe the tracking add-in is of most use in the analysis of heterogenous mixtures of molecules with a rage of diffusivities as our MSD fitting and filtering algorithm cleans up ‘messy’ distributions and allows fitting with a Gaussian mixture model within the GUI allowing for fast and straightforward preliminary analysis. Finally, a basic FRET add-in extends the usefulness of the package for a very common application of single molecule TIRF microscopy.

The package contains two different methods for enhancing the image to allow detection of weak spots. The wavelet transform is computed more rapidly, but the convolution method appears to be more reliable with faint spots in a noisy background, such as might be seen using mCherry as a fluorophore. The choice between these two is left for the user. The automated colocalization procedure to compensate for chromatic aberration is a major practical advantage. However, if colocalization is very low, below the threshold of four spots, then a calibration file will have to be created and used. The display shows the transformation vectors across the screen, which is helpful in reassuring the user that any two spots in different colours are properly related for colocalization.

The improvement in accuracy of step detection was surprisingly large, reaching 20-30% higher than the other methods tested at a signal/noise ratio of 3.5 (Figure 3). Some of the remaining error is the result of close or simultaneous bleaching events that could not be resolved, even though the performance is better than that of a trained observer. This is consistent with the tendency to under-count the number of steps at low signal/noise ratios and the decrease in accuracy as the number of steps increases and the likelihood of events in close proximity increases (Supplementary Figure 24). The results also suggest that any measurement of stoichiometries is likely to be unreliable for three steps or more if the signal/noise ratio is lower than 2.

The measurement of diffusion rates for single molecules presented several difficulties. First, the localization has to be accurate. Again, a signal/noise ratio of >2.5 is helpful. Stage drift needs to be measured and, if sufficiently large, will need to be taken into account. Wobbles or perturbations in focal height should also be avoided. Also of importance is the tuning of the spot-fitting parameters of the program before beginning tracking to ensure that coalescence cannot occur. If two spots coalesce and then separate, the program will not be able to distinguish which particle was which. Setting the maximum jump distance per frame based on the observed motion of the moving particles is also important, especially in fields of view with a high spot density, to ensure valid connections of a particle from one frame to the next. If the value is set too low, the particle will not be tracked as its new position will be treated as a separate object; too high, and the path may jump to a different object.

## 5. Conclusion

We conclude that FluoroTensor is a versatile and useful tool for the analysis of single molecules by wide-field microscopy: its performance in measuring stoichiometry is the best currently available, and innovations such as the automated colocalization and the method for establishing the most reliable measurements of diffusion for each molecule improve the convenience and reliability of methods for analysing the behaviour of single molecules.

### Recommended System Requirements

OS: Windows 10 64 bit

CPU: 8^th^ GEN intel i5-8500 / AMD Ryzen 5 3600XT or newer

RAM: 16GB DDR4 / 32GB DDR4 (recommended for large datasets.)

GPU: (not required) / NVIDIA GTX 1070 (minimum for model training of datasets larger than 1M traces.)

Storage: Minimum 1TB permanent storage for single molecule data (recommend SSD. Use permanent storage for fastest data load times instead of networked storage where possible.)

### Code Availability

We propose to release FluoroTensor alongside this publication (final GITHUB repository link to be confirmed). Video tutorials will be made available on the SpliceSelect website (https://www.spliceselect.org).

## Supporting information

Figures comprising Supplementary Methods and Results

## Acknowledgements

Dr B. Xu (Informetis, Cambridge) provided insights in the early stages of our AI development. Dr A. Revyakin and Dr D. Cherny provided helpful pointers and discussions on enhancement techniques for single molecule images. Dr T.J. Ragan provided insights into data analysis methods. Mr C. Lucas, Dr V. Paschalis and Dr S. Tubasum supplied single molecule TIRF data. Prof. X. Fang and Dr J. Xu kindly provided their CLDNN model for our comparisons.

## CRediT authorship contribution statement

MFKW: Conceptualization, Methodology, Software, Validation, Formal analysis, Writing-original draft, Writing-review and editing, Visualization. CBA: Resources. NH: Methodology, Software, Writing-review and editing. AJH: Conceptualization, Methodology, Writing-review and editing, Supervision. ICE: Conceptualization, Writing-original draft, Writing-review and editing, Supervision, Project administration, Funding acquisition.

## References

1 Leake, M. C. et al. Stoichiometry and turnover in single, functioning membrane protein complexes. Nature 443, 355–358 (2006). https://doi.org:nature05135 [pii] 10.1038/nature05135

2 Shu, D., Zhang, H., Jin, J. & Guo, P. Counting of six pRNAs of phi29 DNA-packaging motor with customized single-molecule dual-view system. EMBO J. 26, 527–537 (2007). https://doi.org:7601506 [pii]10.1038/sj.emboj.7601506

3 Abelson, J. et al. Conformational dynamics of single pre-mRNA molecules during in vitro splicing. Nat Struct Mol Biol 17, 504–512 (2010). 10.1038/nsmb.1767

4 Luo, F., Qin, G., Xia, T. & Fang, X. H. Single-Molecule Imaging of Protein Interactions and Dynamics. Annu Rev Anal Chem 13, 337–361 (2020). 10.1146/annurev-anchem-091619-094308

5 Chen, L. et al. Stoichiometries of U2AF35, U2AF65 and U2 snRNP reveal new early spliceosome assembly pathways. Nucleic Acids Res. 45, 2051-2067 (2017). 10.1093/nar/gkw860

6 Cherny, D. et al. Stoichiometry of a regulatory splicing complex revealed by single-molecule analyses. EMBO J. 29, 2161–2172 (2010). https://doi.org:emboj2010103 [pii] 10.1038/emboj.2010.103

7 Hodson, M. J., Hudson, A. J., Cherny, D. & Eperon, I. C. The transition in spliceosome assembly from complex E to complex A purges surplus U1 snRNPs from alternative splice sites. Nucleic Acids Res. 40, 6850–6862 (2012). 10.1093/nar/gks322

8 Hoskins, A. A., Gelles, J. & Moore, M. J. New insights into the spliceosome by single molecule fluorescence microscopy. Curr Opin Chem Biol 15, 864–870 (2011). https://doi.org:S1367-5931(11)00157-8 [pii] 10.1016/j.cbpa.2011.10.010

9 Jobbins, A. M. et al. Exon-independent recruitment of SRSF1 is mediated by U1 snRNP stem-loop 3. EMBO J. 41, e107640 (2022). 10.15252/embj.2021107640

10 Jobbins, A. M. et al. The mechanisms of a mammalian splicing enhancer. Nucleic Acids Res. 46, 2145–2158 (2018). 10.1093/nar/gky056

11 Ulbrich, M. H. & Isacoff, E. Y. Subunit counting in membrane-bound proteins. Nat Methods 4, 319–321 (2007). https://doi.org:nmeth1024 [pii] 10.1038/nmeth1024

12 Shcherbakova, I. et al. Alternative Spliceosome Assembly Pathways Revealed by Single-Molecule Fluorescence Microscopy. Cell Reports 5, 151–165 (2013). 10.1016/j.celrep.2013.08.026

13 Herbert, K. M. et al. A heterotrimer model of the complete Microprocessor complex revealed by single-molecule subunit counting. RNA 22, 175–183 (2016). 10.1261/rna.054684.115

14 del Rio, A. et al. Stretching single talin rod molecules activates vinculin binding. Science 323, 638–641 (2009). 10.1126/science.1162912

15 Chen, Y., Deffenbaugh, N. C., Anderson, C. T. & Hancock, W. O. Molecular counting by photobleaching in protein complexes with many subunits: best practices and application to the cellulose synthesis complex. Mol Biol Cell 25, 3630–3642 (2014). 10.1091/mbc.E14-06-1146

16 Shepherd, J. W., Higgins, E. J., Wollman, A. J. M. & Leake, M. C. PySTACHIO: Python Single-molecule TrAcking stoiCHiometry Intensity and simulatiOn, a flexible, extensible, beginner-friendly and optimized program for analysis of single-molecule microscopy data. Comput Struct Biotechnol J 19, 4049–4058 (2021). 10.1016/j.csbj.2021.07.004

17 Tsekouras, K., Custer, T. C., Jashnsaz, H., Walter, N. G. & Presse, S. A novel method to accurately locate and count large numbers of steps by photobleaching. Mol Biol Cell 27, 3601–3615 (2016). 10.1091/mbc.E16-06-0404

18 Reyes-Lamothe, R., Sherratt, D. J. & Leake, M. C. Stoichiometry and architecture of active DNA replication machinery in Escherichia coli. Science 328, 498–501 (2010). 10.1126/science.1185757

19 Wakelin, S. & Bagshaw, C. R. A prism combination for near isotropic fluorescence excitation by total internal reflection. J Microsc 209, 143–148 (2003).

20 Messina, T. C., Kim, H., Giurleo, J. T. & Talaga, D. S. Hidden Markov model analysis of multichromophore photobleaching. J Phys Chem B 110, 16366–16376 (2006). 10.1021/jp063367k

21 McGuire, H., Aurousseau, M. R., Bowie, D. & Blunck, R. Automating single subunit counting of membrane proteins in mammalian cells. J Biol Chem 287, 35912–35921 (2012). 10.1074/jbc.M112.402057

22 White, D. S., Goldschen-Ohm, M. P., Goldsmith, R. H. & Chanda, B. Top-down machine learning approach for high-throughput single-molecule analysis. Elife 9 (2020). 10.7554/eLife.53357

23 Xu, J. et al. Automated Stoichiometry Analysis of Single-Molecule Fluorescence Imaging Traces via Deep Learning. J Am Chem Soc 141, 6976–6985 (2019). 10.1021/jacs.9b00688

24 Wan, L., Zeiler, M., Zhang, S., Le Cun, Y. & Fergus, R. in International conference on machine learning. 1058–1066 (PMLR).

25 Srivastava, N., Hinton, G., Krizhevsky, A., Sutskever, I. & Salakhutdinov, R. Dropout: A Simple Way to Prevent Neural Networks from Overfitting. J Mach Learn Res 15, 1929–1958 (2014).

26 Mysková, J. et al. Directionality of light absorption and emission in representative fluorescent proteins. Proc Natl Acad Sci USA 117, 32395–32401 (2020). 10.1073/pnas.2017379117

27 Michalet, X., Pinaud, F. & Weiss, S. Single Quantum Dot Trajectory Analysis: Beyond the Single Diffusion Mode Model. Biophys J 98, 203a–204a (2010). DOI 10.1016/j.bpj.2009.12.1086

28 Young, G. et al. Quantitative mass imaging of single biological macromolecules. Science 360, 423–427 (2018). 10.1126/science.aar5839

29 Chen, J. & Gupta, A. K. Testing and locating variance changepoints with application to stock prices. J Am Stat Assoc 92, 739–747 (1997). Doi 10.2307/2965722

30 Marguet, D., Lenne, P. F., Rigneault, H. & He, H. T. Dynamics in the plasma membrane: how to combine fluidity and order. EMBO J. 25, 3446–3457 (2006). DOI 10.1038/sj.emboj.7601204

31 Ruthardt, N., Lamb, D. C. & Bräuchle, C. Single-particle Tracking as a Quantitative Microscopy-based Approach to Unravel Cell Entry Mechanisms of Viruses and Pharmaceutical Nanoparticles. Molecular Therapy 19, 1199–1211 (2011). 10.1038/mt.2011.102

32 Zalejski, J., Sun, J. C. & Sharma, A. Unravelling the Mystery inside Cells by Using Single-Molecule Fluorescence Imaging. J Imaging 9 (2023). https://doi.org:ARTN 192 10.3390/jimaging9090192

