## Supplementary material for "FluoroTensor: identification and tracking of colocalised molecules and their stoichiometries in multi-colour single molecule imaging via deep learning": Figures comprising Supplementary Methods and Results

### FluoroTensor Supplementary Information

#### Section 1. Supplementary Methods

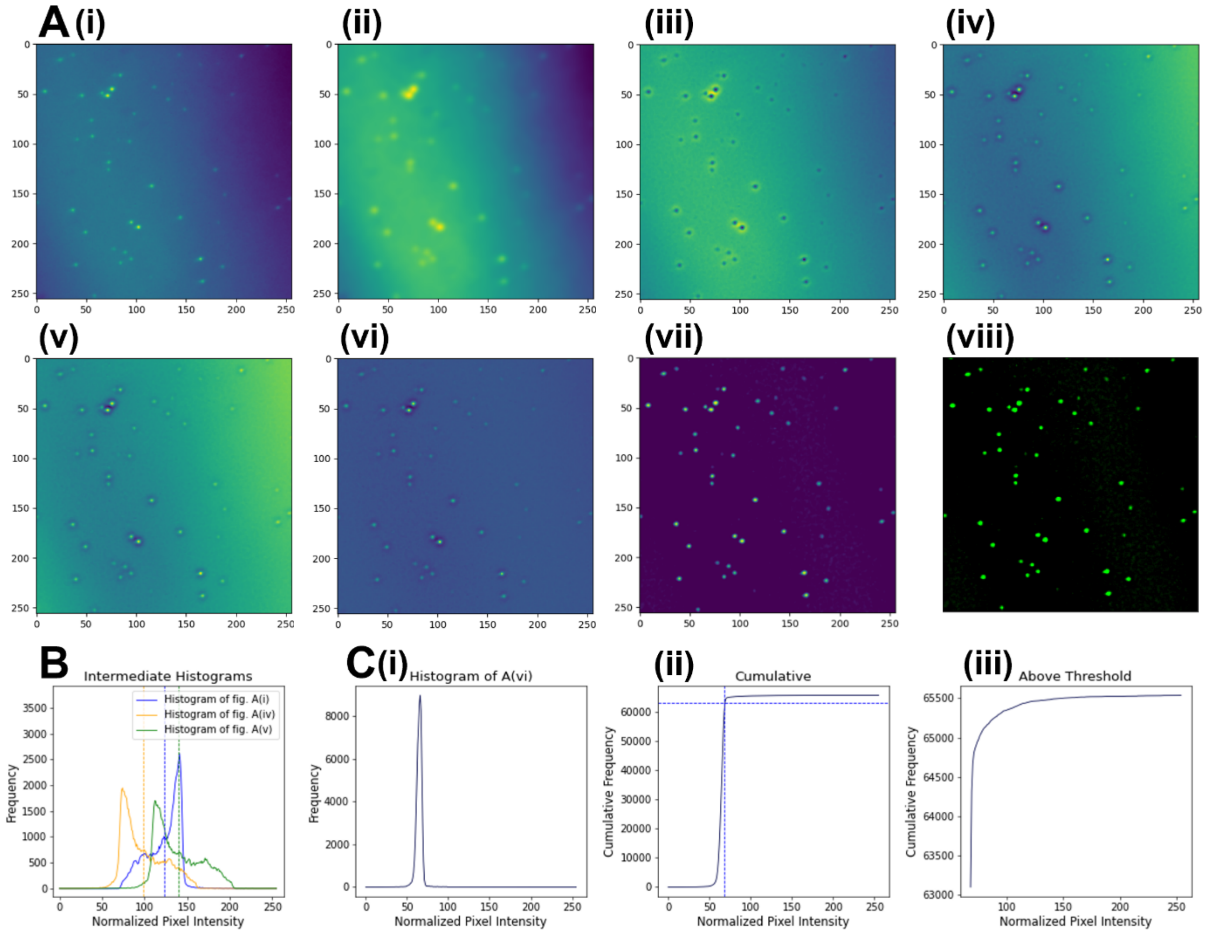


**Figure 1. (A)** Stages of the image enhancement process. **(i)** The original image. **(ii)** The convolution of (i) with a Gaussian kernel. **(iii)** The residual of (ii) – (i). **(iv)** Inverted form of (iii). **(v)** Nonlinear mapping applied to (iv). **(vi)** Result of multiplicative merging of (i) and (v). **(vii)** Image (vi) after background subtraction. **(viii)** False colour version of (vii) displayed in the program. **(B)** Histograms of the original image (A)(i) in blue, inverted residual (A)(iv) in yellow, and nonlinear mapped form (A)(v) in green. Raising (A)(iv) to the power calculated using supplementary equation 1 to map (A)(iv) onto (A)(v) scales the negative autofluorescence profile such that it cancels out with the original to produce the uniform background in (vi). **(C)** Thresholding method to subtract the uniform background in (A)(vi) to obtain (A)(vii). **(i)** The histogram of A(vi). **(ii)** The cumulative plot of the histogram in (i). The threshold is set to keep pixel intensities at the 96$^{th}$ percentile and above. **(iii)** Shows the portion of the cumulative pixel intensities above the threshold which should contain the fluorescent foci sans the background. Images were normalized between every step of this enhancement.

$$\mathrm{POWER} = \frac{\log(0.55)}{\log(\mathrm{MEAN\_INTENSITY})} \quad \text{[Supplementary Eq. 1]}$$

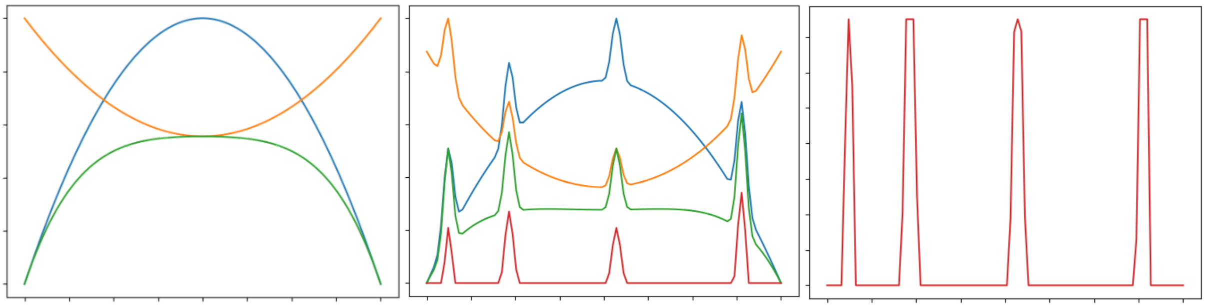

**Figure 2**. Consider this exaggerated representation of a 1-pixel wide slice across the centre of the original image and the inverted convolution residual represented by a 1-dimensional plot where the x axis is the x coordinate of the pixels and the y axis the brightness. **Left**: The blue parabola is a loose representation of the profile of the autofluorescence background across the original image and the orange parabola represents the same for the inverted convolution residual. Note that they both exist on a normalized scale and that owing to their different shapes, the product of the two shown in green still varies significantly from the edges to the centre. Note that the non-uniformity of the background has been somewhat exaggerated to illustrate the point. **Centre**: the original image is represented in blue, now with peaks where the 1-dimensional slice crosses spots. Shown in orange, the inverted convolution residual has had a non-linear mapping applied to it in the form of raising every value to a certain power and renormalizing it, the effect of which scales the parabola such that it cancels out the original autofluorescence while reinforcing the spots to a sufficient extent that a linear subtraction will remove the background almost altogether without thresholding away the spots – shown in green. In red, the slice of the image with only the spots after the background has been linearly subtracted. **Right**: After linear background subtraction the spots are clipped to increase the contrast and renormalized.

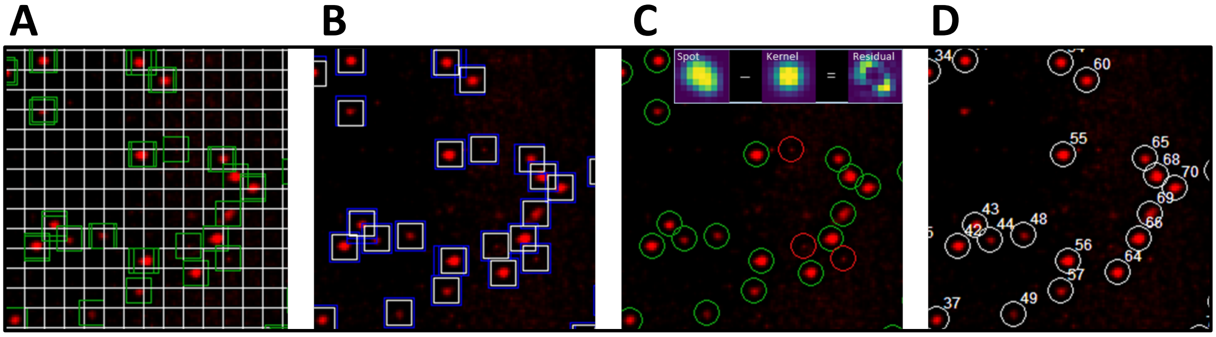


**Figure 3.** Different stages of the spot detection algorithm are shown for a small area in the imaging field. **(A)** The field of view is split into 8x8 pixel grids. Green boxes are centred on the weighted average position of any aberrantly bright object in the image once per 8x8 box. Multiple green boxes around the same object indicate it lies on the boundary between 2 or more 8x8 boxes. **(B)** If multiple instances are detected they are averaged together if within 2 pixels. The bounding box is then re-centred around the spot as the weighted average of the pixel positions. **(C)** The residual between the spot and a pre-computed gaussian kernel is used as a 'wheat from chaff' filter to separate likely potential spots (in green) from aberrant bright regions such as salt and pepper noise or dead pixels (in red). **(D)** The spots that pass the kernel residual threshold are fitted with a 2D Gaussian function with further thresholds based on the Gaussian parameters and to obtain the coordinates with sub-pixel precision.

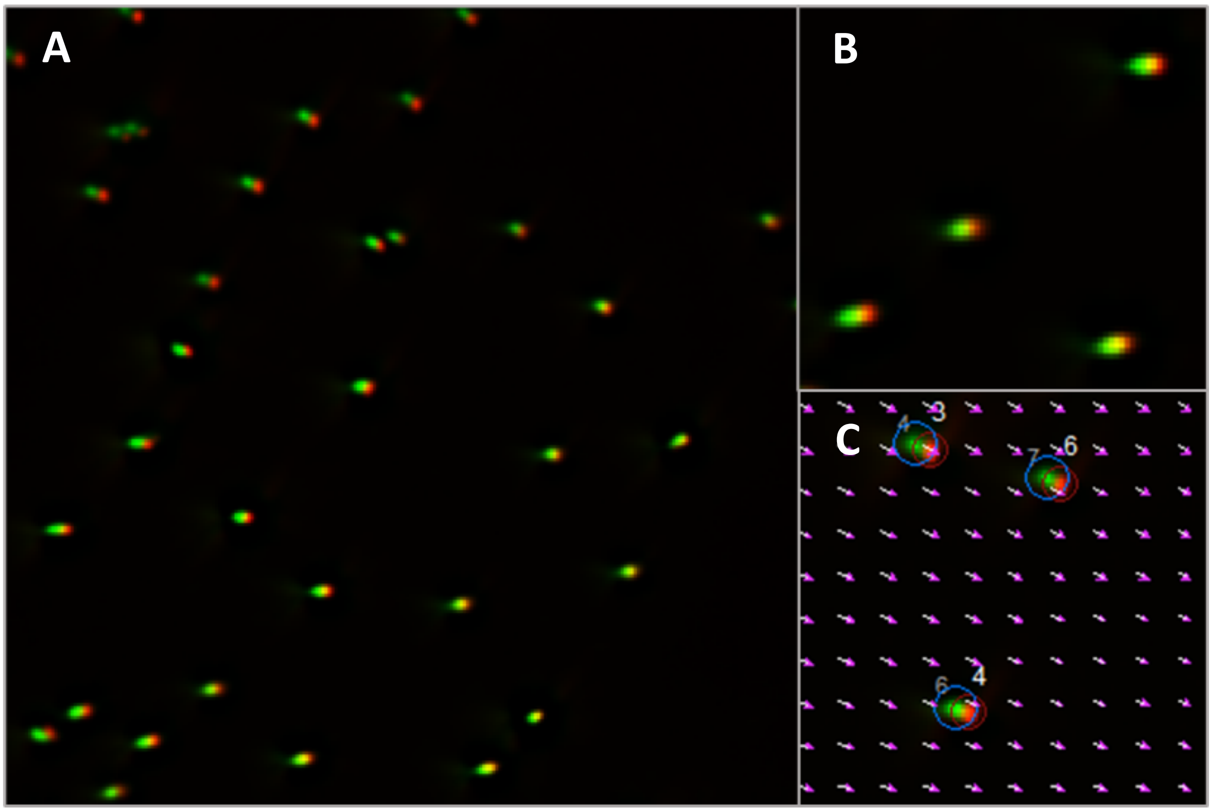


**Figure 4**. Chromatic aberration is seen as a shift in positions from one channel to the next when the images are superimposed. **(A)** Composite image of fluorescent beads excited with a red laser (640nm, shown in red) and the same beads excited with a blue laser (488nm, shown in green) The origin of chromatic aberration (where spots perfectly superimpose) is in the lower right-hand side of the image. **(B)** Enlarged region of the lower left of (A) shows stretched spots in 488nm channel due to the broad emission spectrum of the beads having a range of dispersions. **(C)** The vector field calculated using formulae 1a and 1b in the main paper showing the transformation needed to map the coordinates of spots in the blue channel (488nm, mEGFP) to spots in the red channel (640nm, Cy5). (Note: here we refer to colour channels by excitation laser wavelengths, the transform was calculated using peak emission wavelengths of Cy5, mCherry and mEGFP.)

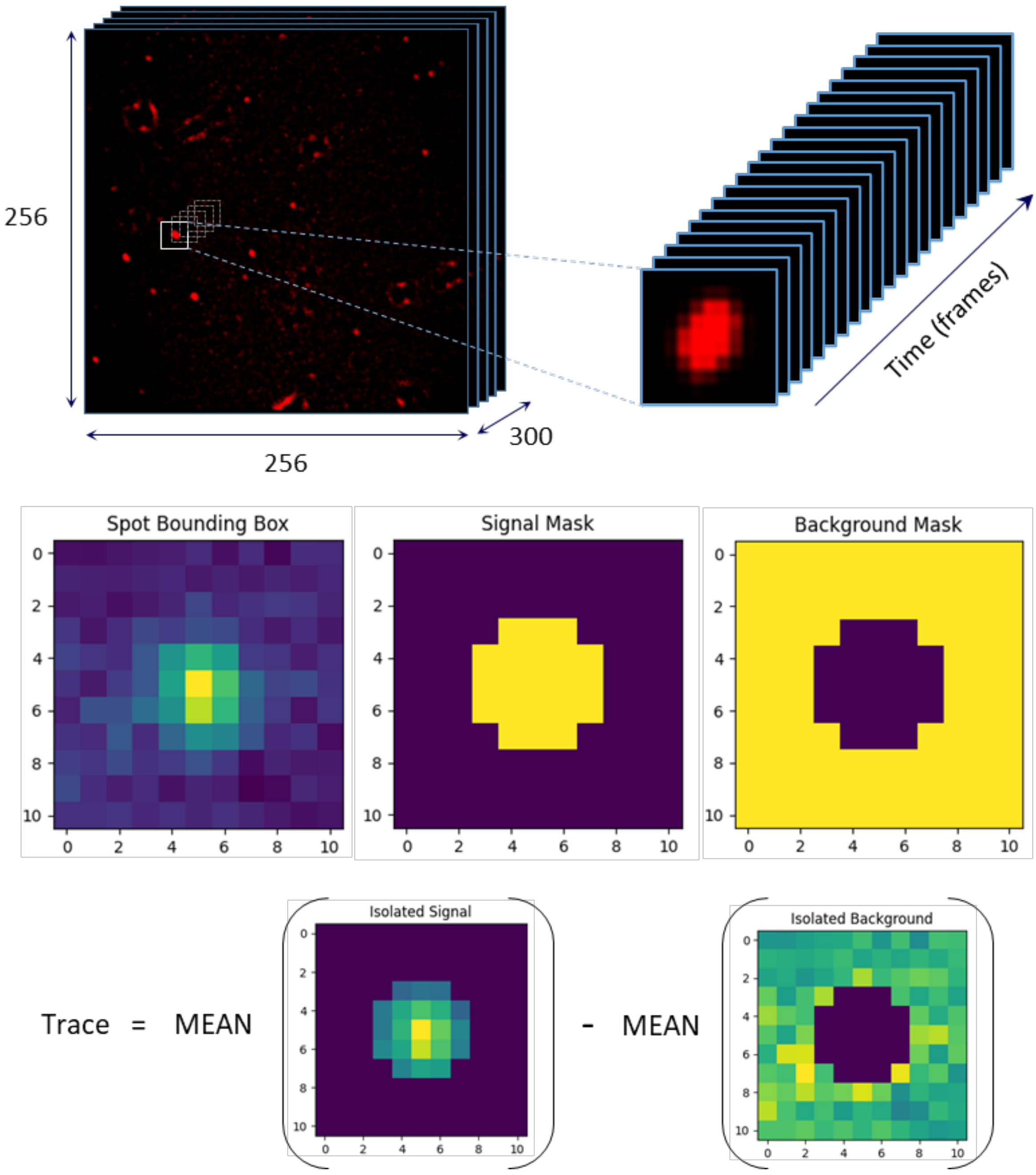


**Figure 5**. The 3D array the movie is stored in is sliced to produce a bounding volume around the spot in with dimensions 11x11xN where N is the number of frames (Our standard operating procedure is to record for 300 frames). For each frame, the 11x11 pixel bounding box around the spot is masked to Isolate the signal and the background. The mean background is then subtracted from the mean signal to obtain the background subtracted intensity. This process produces a fluorescence intensity bleach that reaches a mean intensity of zero when all fluorophores have bleached, allowing for the ability of the neural network to detect incompletely bleached complexes.

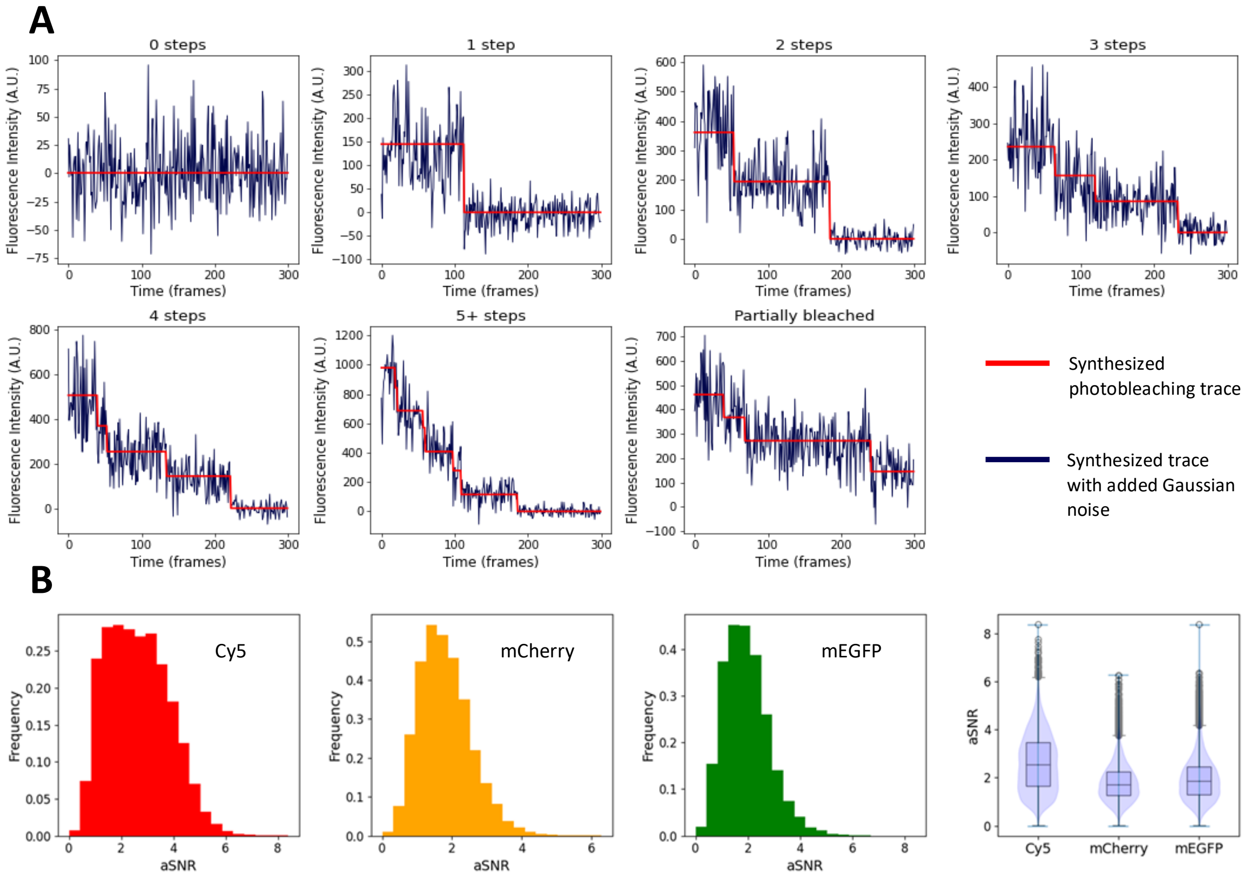


**Figure 6. (A)** Synthesized Cy5 traces for each classification 0-4 steps, 5 or more, and partially bleached (when not all fluorophores photo-bleach within the recording time and the stoichiometry is ambiguous). **(B)** Distributions of signal to noise ratios of the simulated traces used to train the CRNN.

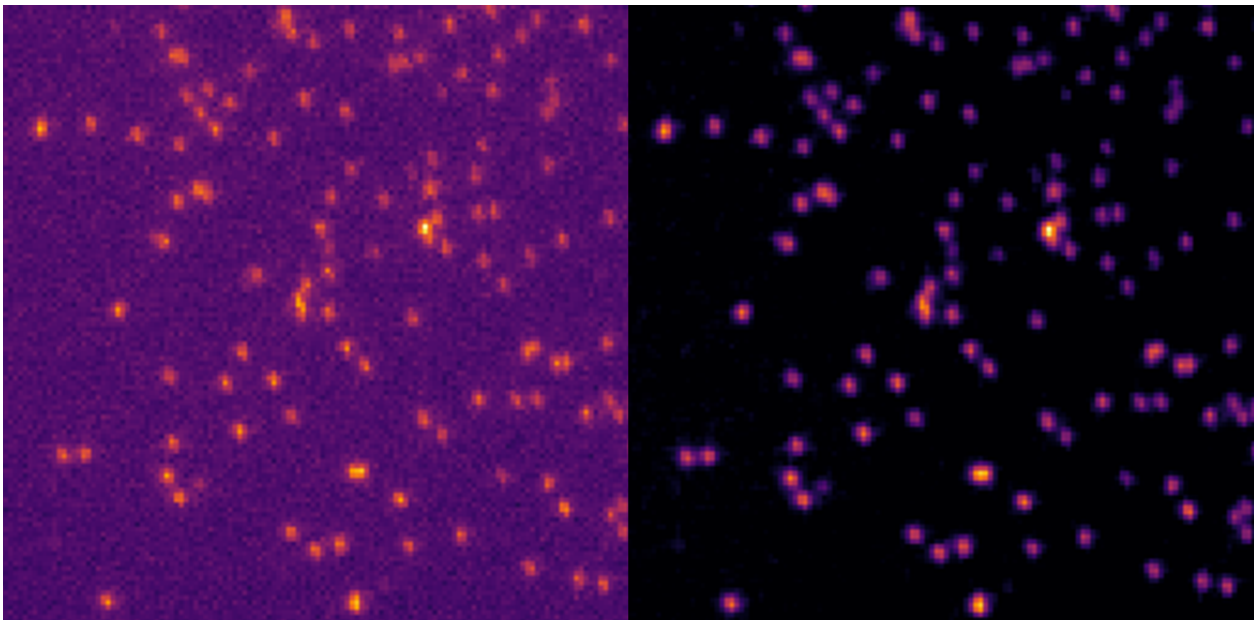

**Figure 7. Left**: A region of a frame from a single molecule movie. **Right**: The same frame after the high pass filtering method was applied to enhance the frame.

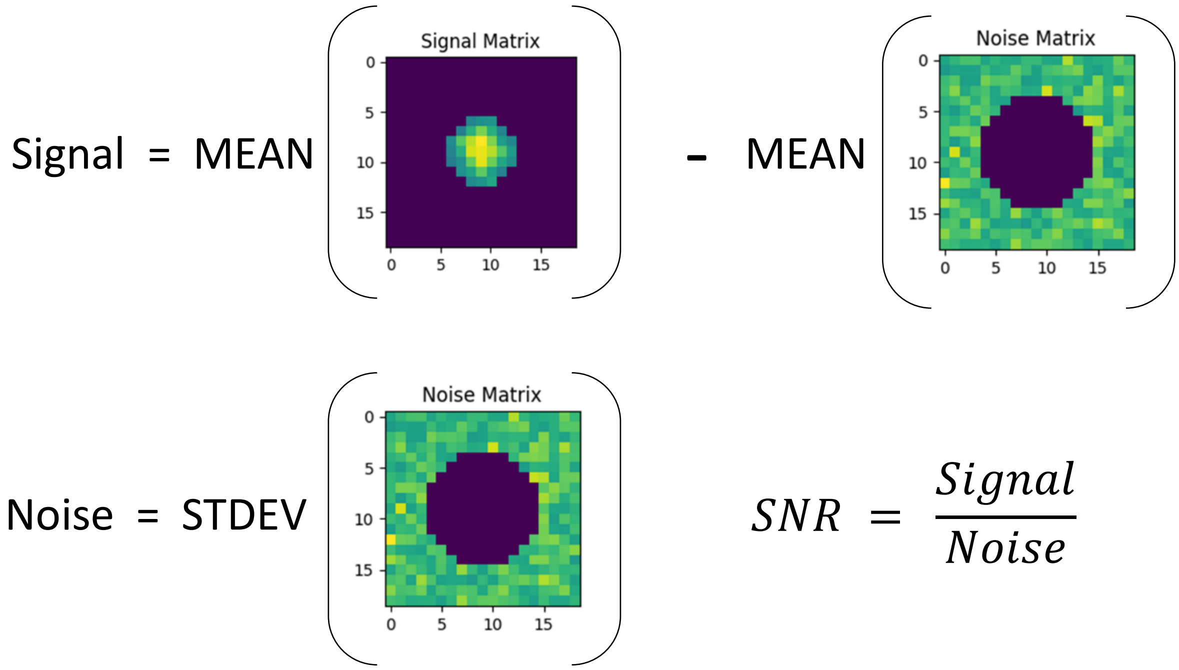


**Figure 8.** Method for calculating the signal to noise ratio (SNR) of a fluorescent molecule in a single frame from an SM movie. First, all frames are cropped around the spot keeping the cropped region centred on the spot taking care that no other spots appear in this region. The cropped frames are summed to reduce noise so the spot can be fitted with a gaussian. This fit returns the full width half at half maximum which is then used to create a mask to isolate the signal from the noise. The background subtracted mean of the signal is divided by the standard deviation of the noise to obtain the SNR for each frame. The SNR for all frames is averaged to obtain the mean signal to noise ratio of the fluorescent molecule over the time it is tracked.

#### Section 2. Supplementary Results

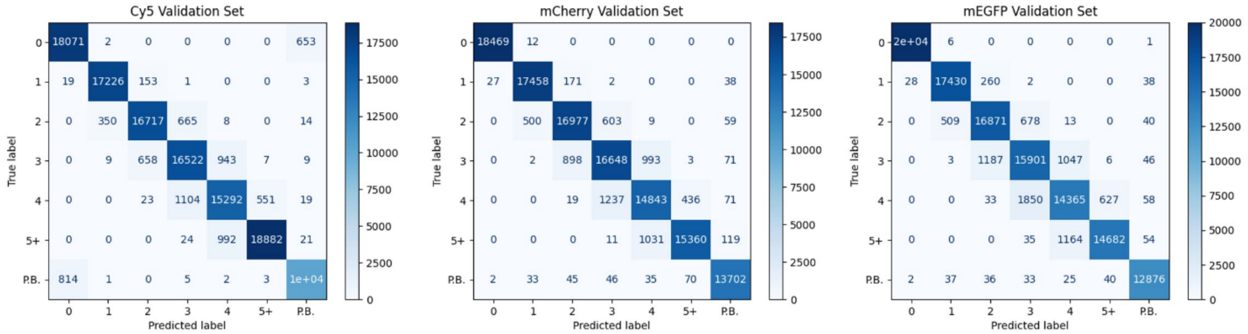


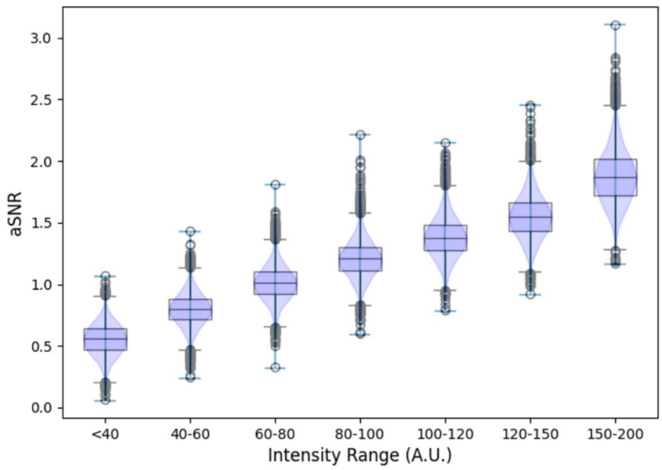


| Arbitrary Intensity Range | Minimum Plateau Length | Mean aSNR |
|---|---|---|
| Int ≤ 40 | 23 | <0.557 |
| 40 < Int ≤ 60 | 18 | 0.802 |
| 60 < Int ≤ 80 | 12 | 1.012 |
| 80 < Int ≤ 100 | 8 | 1.210 |
| 100 < Int ≤ 120 | 5 | 1.383 |
| 120 < Int ≤ 150 | 3 | 1.550 |
| 150 < Int ≤ 200 | 2 | 1.877 |
| Int > 200 | 1 | N/A |

**Figure 9. Top**: Validation set confusion matrices. Each validation set contained 120,000 traces with distributions of intensities, bleaching times, and all other attributes mirroring the training set. These sets were only seen by the model in the inference step of each training epoch to determine if the validation accuracy had increased from the previous epoch to determine if the model would be saved. This system was implemented as a form of early stopping to prevent overfitting. **Bottom**: Signal to noise ratio distributions for different ranges of arbitrary intensities of simulated fluorophores and how these intensities are used to set a minimum plateau length restriction.

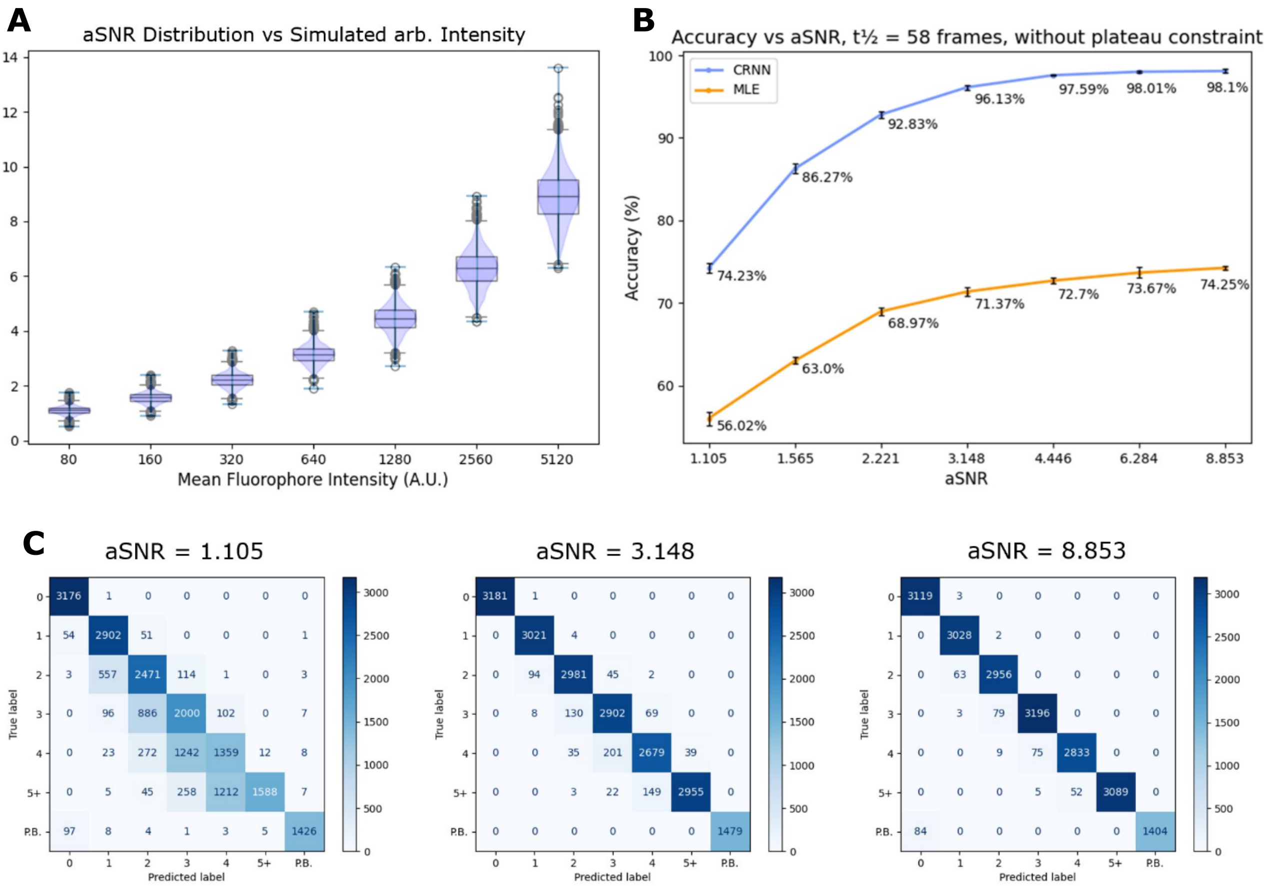


**Figure 10. (A)** Signal to noise ratio distribution of datasets generated with different arbitrary intensities of simulated fluorophores. **(B)** Accuracy of the model vs signal to noise ratios. The bleaching rate is represented by the half-life of the fluorophores calculated as follows: $t_{\frac{1}{2}} = \frac{ln\,(2)}{P(B)}$ where P(B) is the probability a fluorophore will bleach in any frame. For this accuracy test $t_{\frac{1}{2}} = 58$ frames. **(C)** Confusion matrices for datasets with aSNRs 1.105, 3.148, and 8.853 respectively. Note that at very low signal to noise ratios, errors are due to under predictions of the ground truth likely due to plateaus hidden in the noise.

### Figure 11 part 1

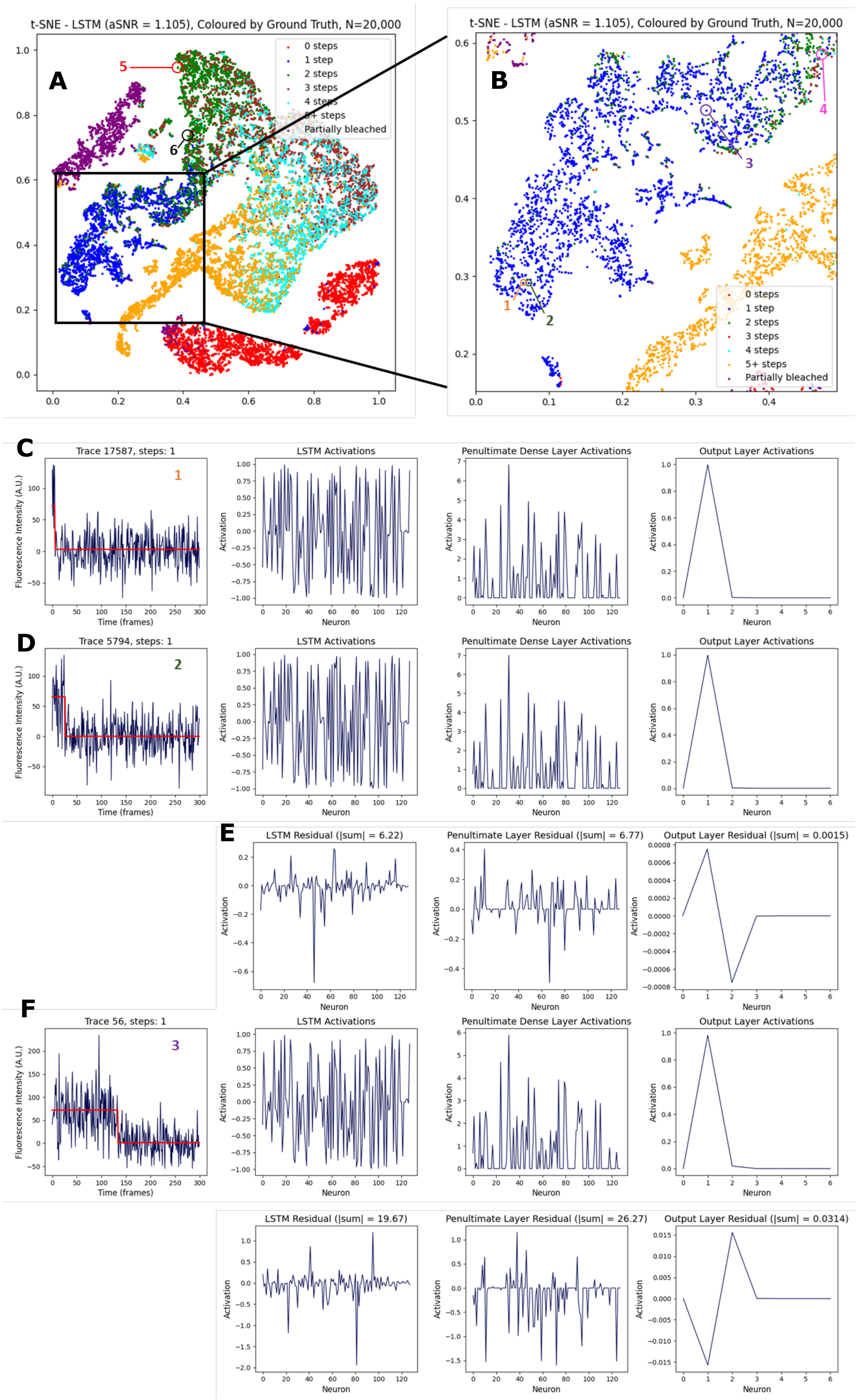

#### Figure 11 part 2

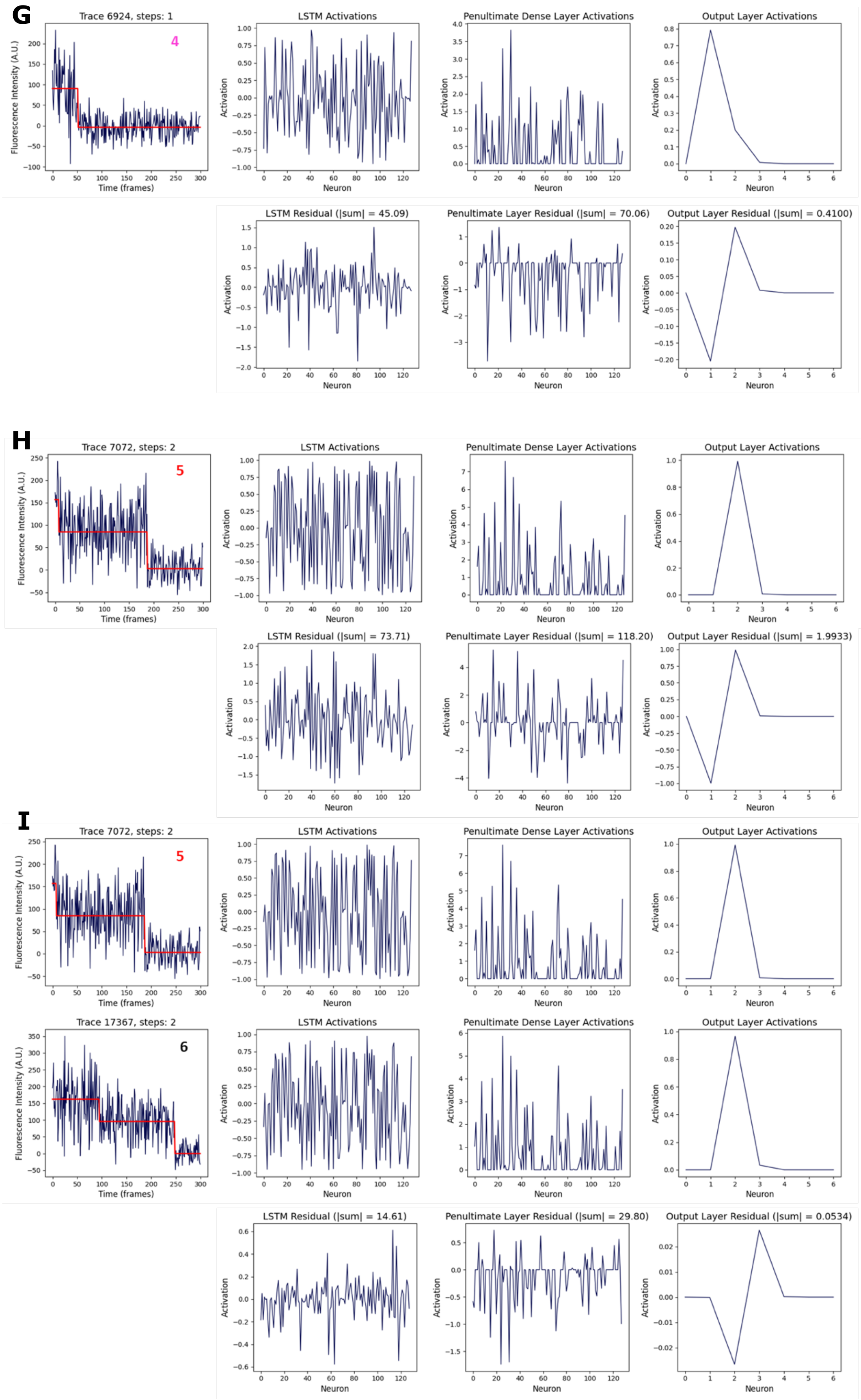

**Figure 11. (A)** A t-SNE plot of the activations of the LSTM layer of the model on the dataset of 20,000 traces from figure 8 coloured by ground truth. **(B)** A closeup of the 1 step cluster. Certain points on this plot were cross-referenced back to the original trace; these points have been numbered. **(C)** The original trace for point 1 is shown, to the right of the trace are the LSTM feature vector, the penultimate dense layer activations and finally the output activations respectively. **(D)** The same plots for a very nearby point in the cluster. Note that both traces have a single clear step and thus belong in the same classification even though the step occurs at a different frame and every single intensity value of the noise will vary greatly. Despite these significant differences of values from corresponding time points of the two traces, the feature vectors extracted by CNN / LSTM layers are extremely similar. **(E)** The corresponding residuals calculated by subtracting the respective activations of trace 2 from trace 1. The residuals are small and the magnitude of the absolute sum of the residuals are small. **(F)** The same plots for trace 3 and the residuals between 1 and 3. As expected the residuals are slightly larger for traces separated by greater distances in the t-SNE plot as such distances are an abstract representation of similarity. **(G)** This same comparison has been made for trace 4 which is at the upper right corner of the 1-step cluster near the overlap to the 2-step cluster. The noise in the first plateau appears to fluctuate such that an ambiguous looking plateau appears within it with a lower mean. This was verified by attempting to fit the trace with 2 steps even though we know from the ground truth that the trace only has one virtual fluorophore. The model still classifies this trace as 1 step correctly but note the high activation of the neuron for 2 steps to the right due to the ambiguity. As expected, the residual between trace 1 with a single clear step and trace 4 is much higher than the other 1-step traces and the activation pattern neither resembles 1 step or 2 steps but a mixture of activations between both classes. **(H)** The same comparison has been made between trace 1 and trace 5, a trace with a ground truth of 2 steps. The activation patterns are distinctly different and the residuals are much larger. **(I)** a final comparison between two 2-step traces, namely traces 5 and 6 which both have 2 steps but completely different intensity profiles and step positions. Once again, the model has extracted the features and produced very similar feature vectors for these traces which are both decoded by the dense layers to give output activations strongly in favour of assigning 2 steps.

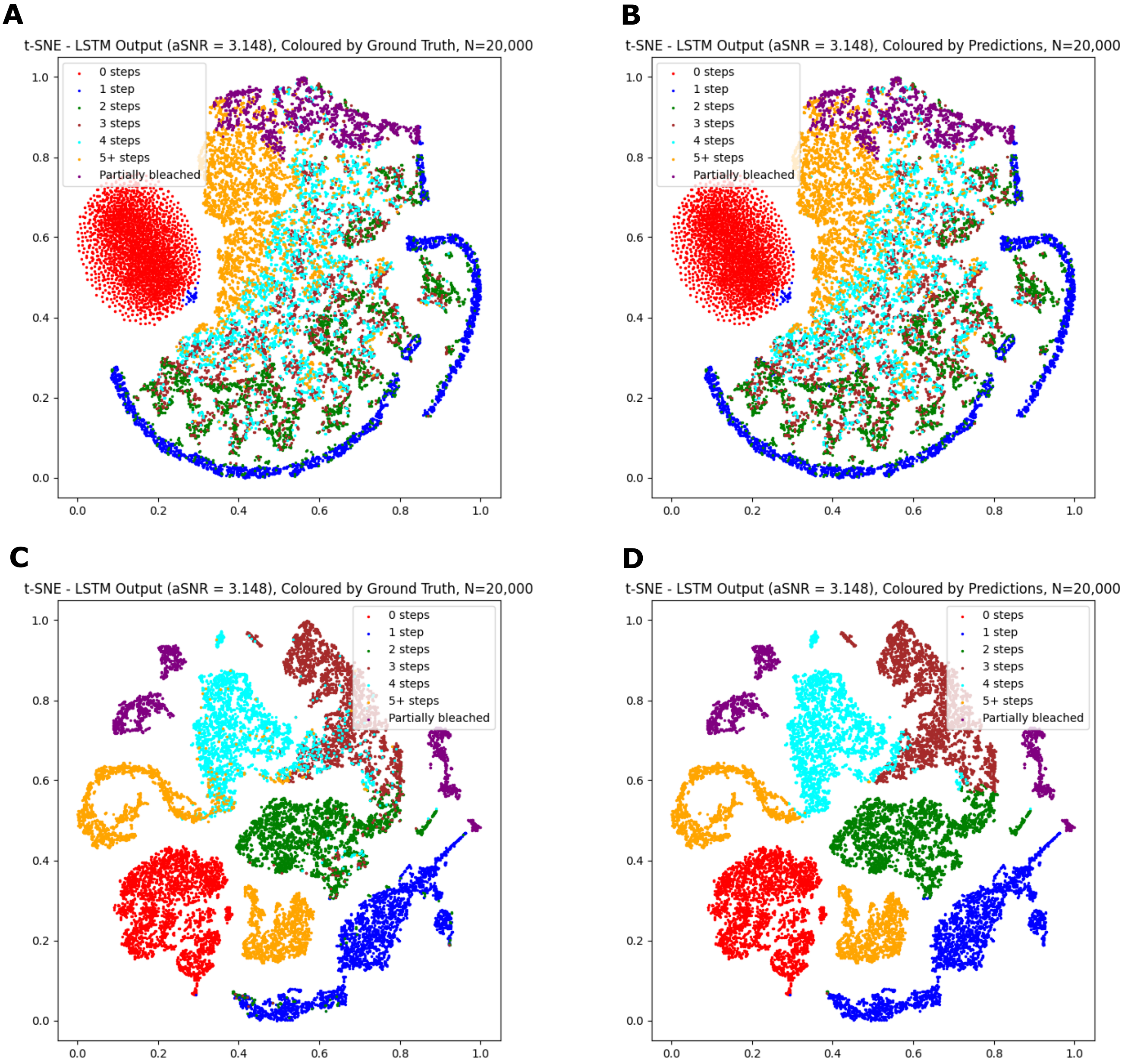


**Figure 12. (A)** t-SNE of 20K traces of mean aSNR = 3.148 coloured by ground truth. **(B)** Same as (A) but coloured by predictions made by our neural network model on the traces. **(C)** t-SNE of LSTM output feature vectors showing clear clustering of traces with the same step count. **(D)** Same as (C) but coloured by neural network predictions.

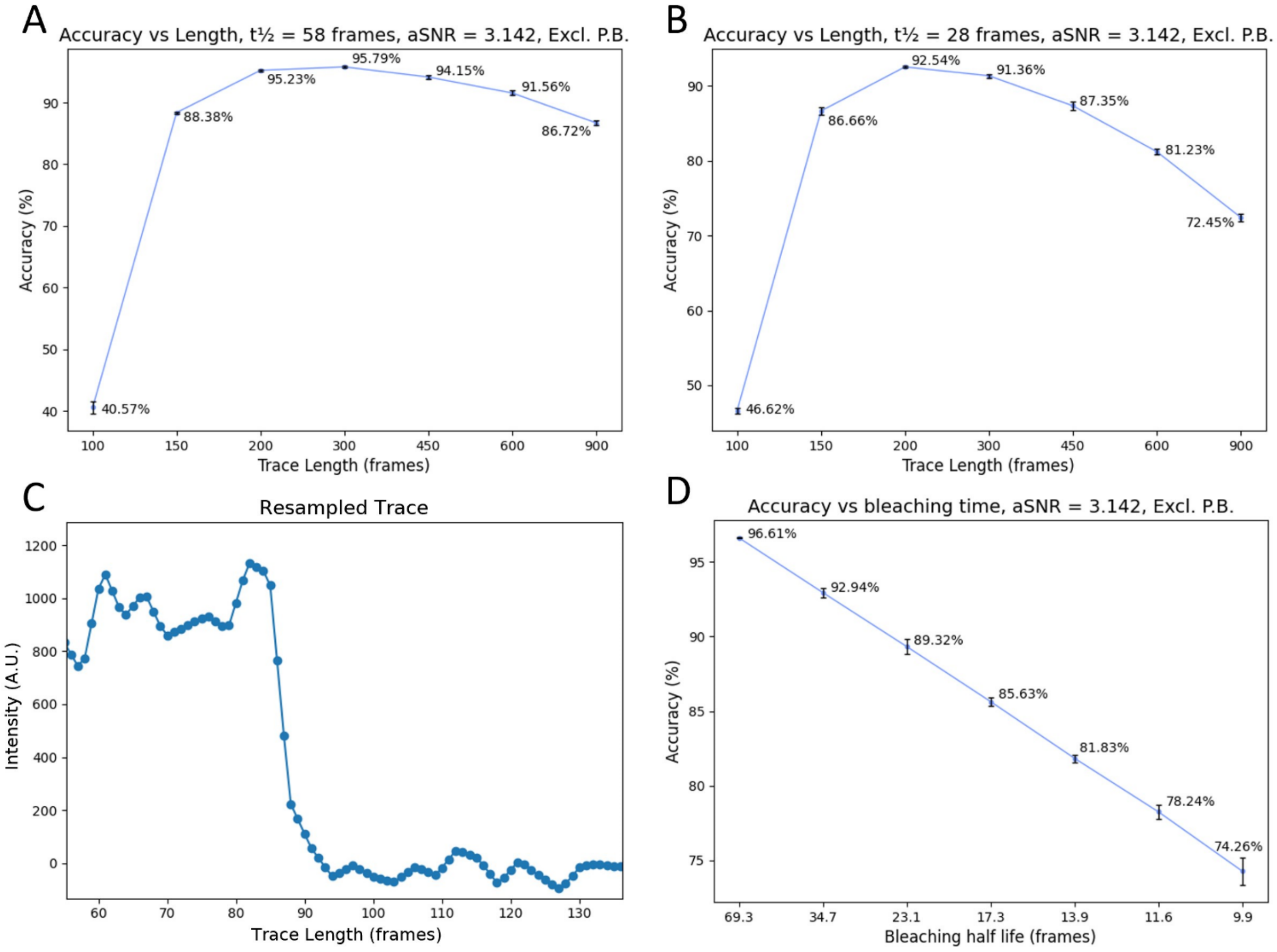


**Figure 13.** An investigation into factors that affect prediction accuracy. **(A)** Accuracy of model when predicting steps in datasets of resampled traces vs original trace length. aSNR = 3.142, photo-bleaching half-life = 58 frames. **(B)** Same as (A) but photo-bleaching half-life = 28 frames. **(C)** Close-up of a step in a 100-frame trace resampled up to 300 frames using a linear interpolator. **(D)** Accuracy of predictions vs photo-bleaching half-life. Half-life values from left to right are equal to probabilities: 1%, 2%, 3%, 4%, 5%, 6%, 7%, of a fluorophore to bleach per frame respectively. In all figures (A) – (D), Partially bleached traces were removed from the datasets as different trace lengths and bleaching rates vary the representations of this class and as the network is exceptional at detecting partially bleached traces, this would skew the accuracy percentages.

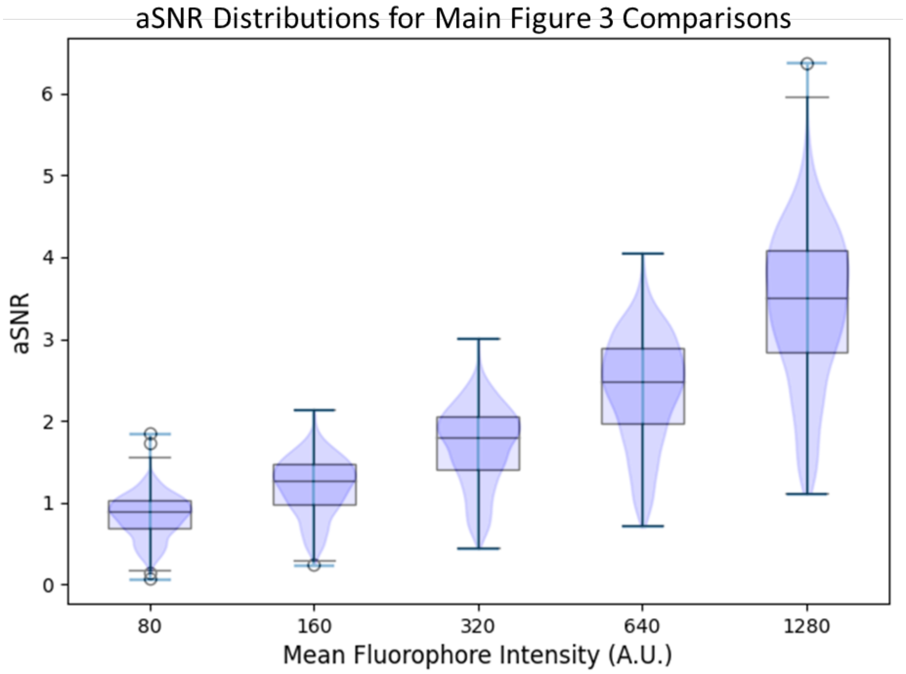


**Figure 14.** aSNR distribution of synthetic fluorescent protein-like traces used for comparisons between FluoroTensor, CLDNN and MLE in main figure 3.

$$t_{\frac{1}{2}} = \frac{\ln(2)}{\frac{P}{n}\int_{n}^{0}\cos^2 x\,dx}$$ [Supplementary Eq. 2]

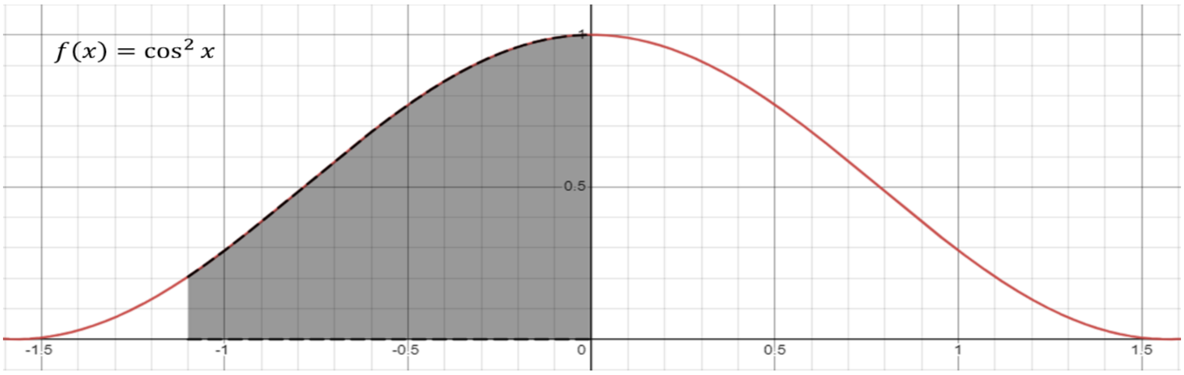


**Figure 15.** Fluorescent proteins are assumed to adsorb onto glass slides at random orientations. The distribution of orientations that allow for efficient absorption of photons and orthogonal radial distribution of emitted photons that follow a path through emission pathway optics to reach the EMCCD chip at these orientations give rise to a $\cos^2$ distribution of intensities. Our simulations follow this distribution with a linear relationship between intensity (photons emitted) and framewise photobleaching probabilities. Only the region of the distribution shown in grey was used since the low intensity fluorophores at the fringes of the distribution are unlikely to be detected at the image processing stage. Supplementary equation 2 was used to calculate the mean half life where P is the maximum possible simulated bleaching rate (at x=0 of the $\cos^2$ distribution) and 1/n * the integral gives the mean bleaching rate of the distribution where n is the lower limit of integration (the low intensity cut-off).

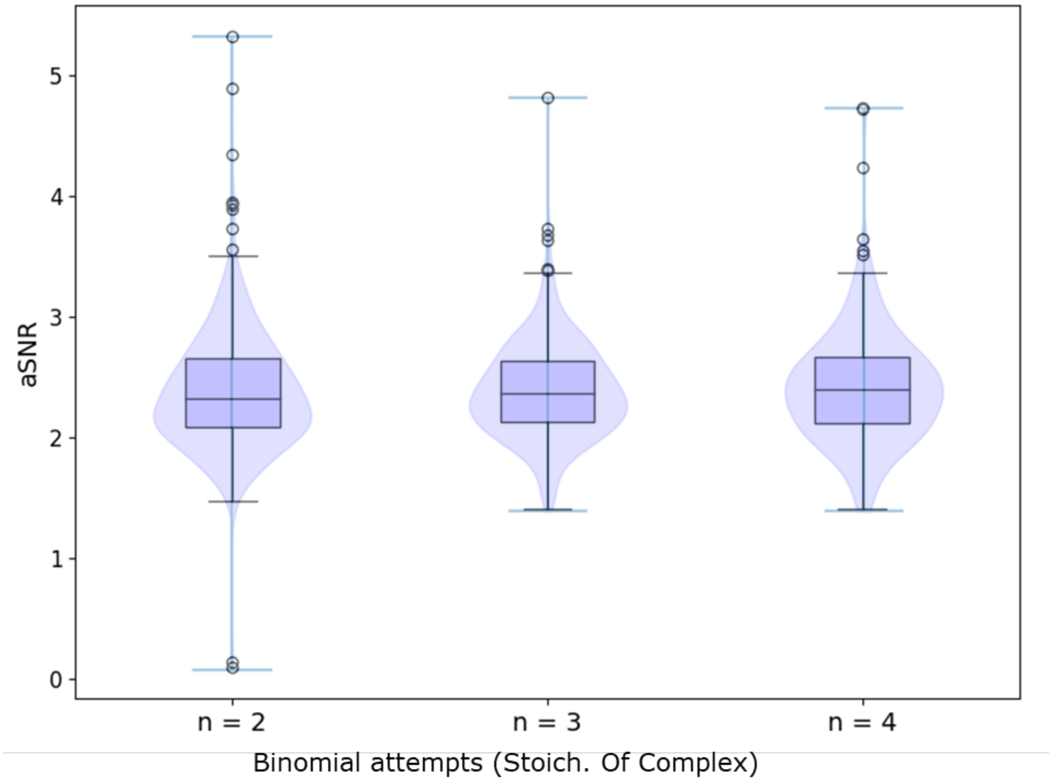


**Figure 16.** aSNR distribution of traces in each simulated TIF stack analysed and shown in main figure 4 for each dataset with a known stoichiometry.

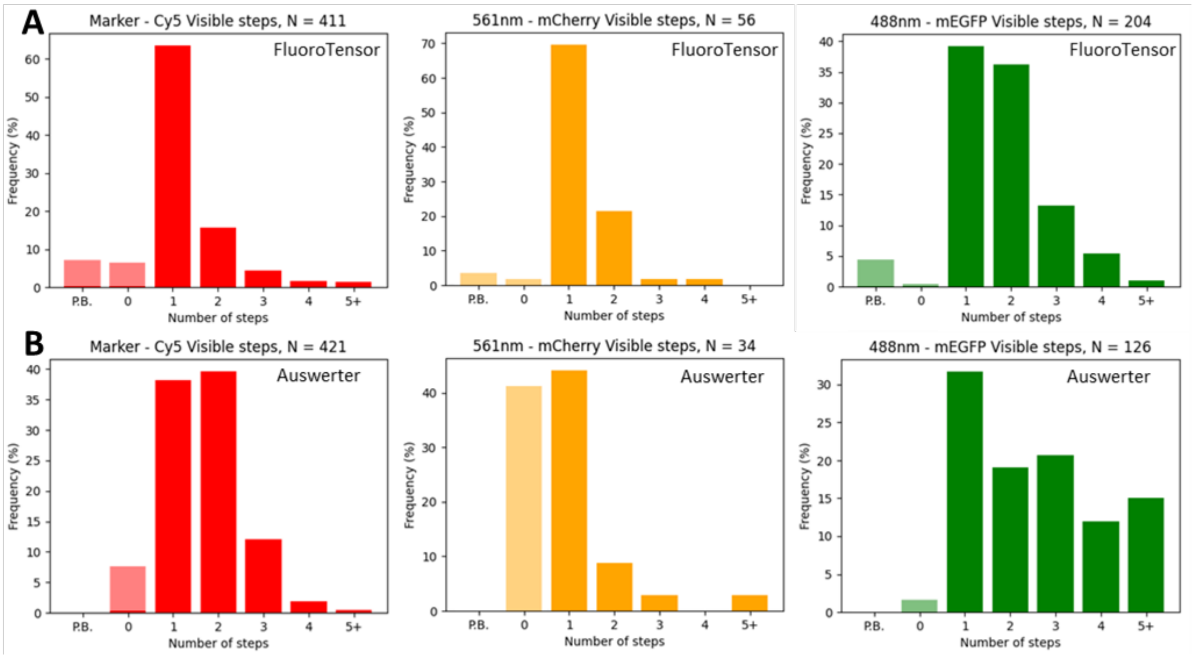


**Figure 17. (A)** Colocalization experiment analysed with FluoroTensor. Step distributions are shown for colocalized spots of mCherry labelled U1A and mEGFP labelled SRSF1 on a Cy5-labelled RNA substrate containing 3 SRSF1 binding motifs {Jobbins, 2022 #65}. **(B)** The same experiment as in (A) but analysed manually in a MATLAB suite (Auswerter). For both (A) and (B) the protein distributions are shown for spots colocalized with a single step Cy5 trace (i.e., a single pre-mRNA).

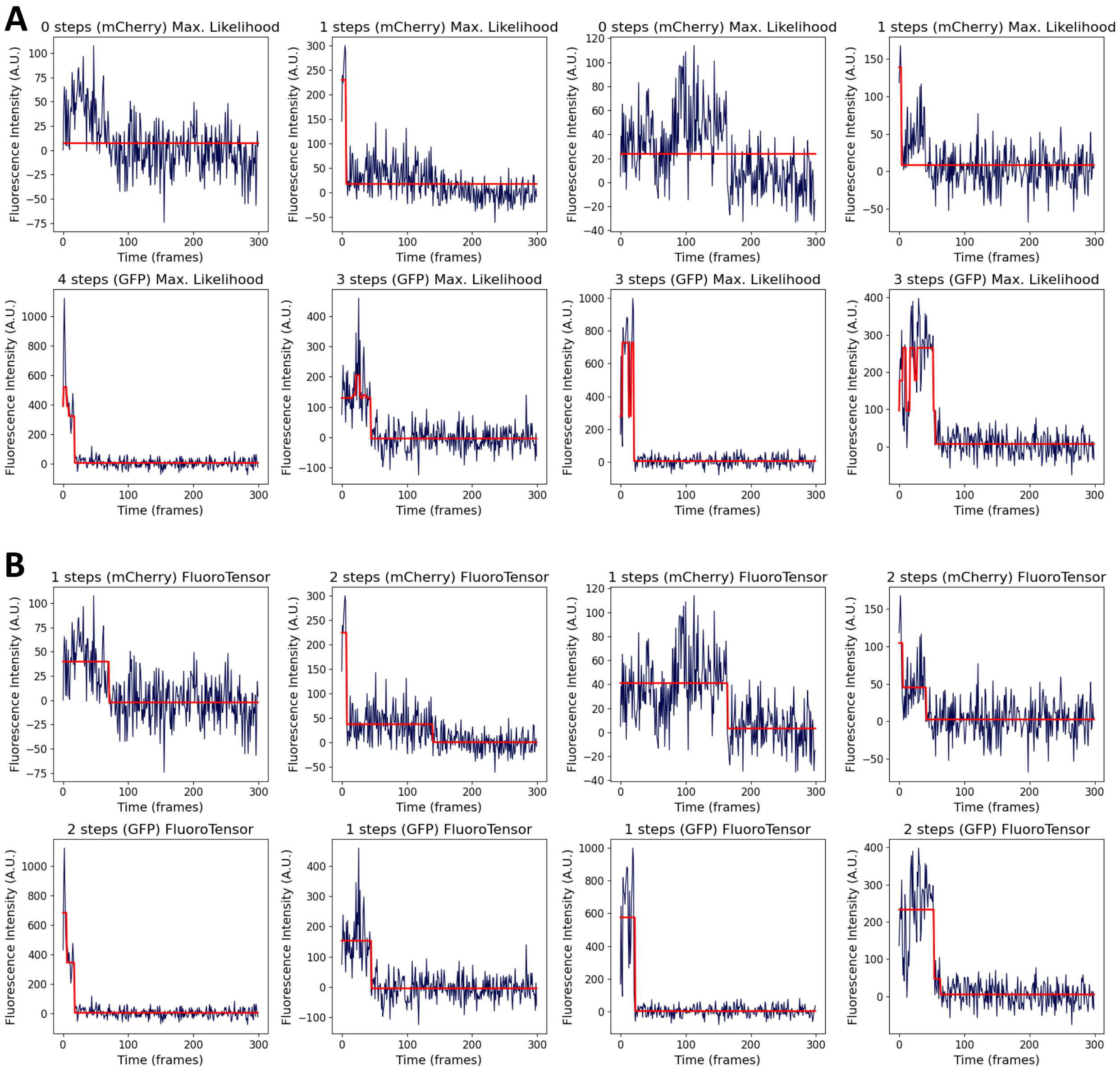


**Figure 18. (A)** Examples of errors in step detection in the Auswerter MATLAB program using a maximum likelihood estimator. Top row shows examples of a consistent undercounting of mCherry steps when the minimum step-size threshold is too low. The next row below that shows the problems that occur when reducing the threshold: examples of GFP traces where the system is finding more steps than it should. **(B)** The exact same traces with steps predicted by the neural network models in FluoroTensor. The red fit lines for the traces analysed by FluoroTensor were fitted manually once the step counts were predicted by the models. FluoroTensor has an automated trace fitting algorithm that uses the step count from the model as an input and uses a moving average window to find the n largest drops in intensity where n is the number of steps however this system is entirely separate from the model used to find the number of steps.

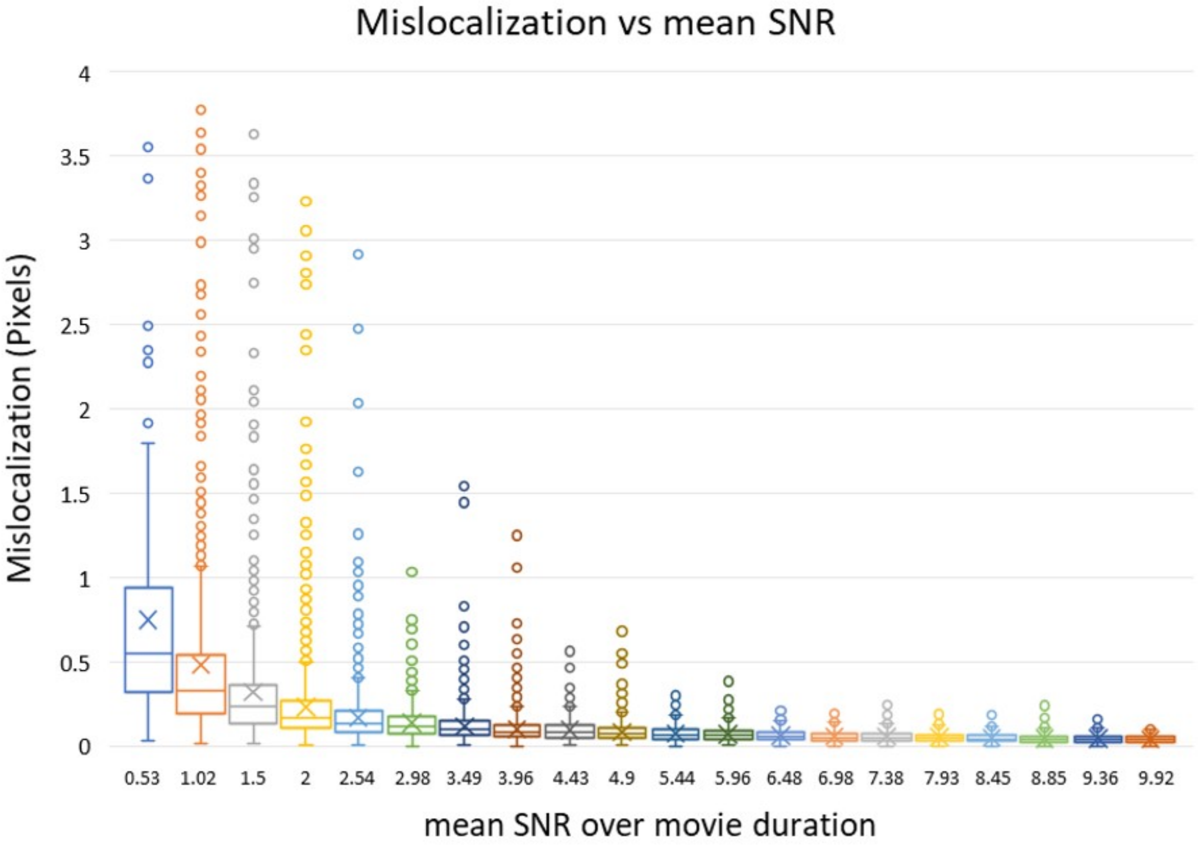


**Figure 19.** Distributions of mislocalization distances of a simulated stationary fluorescent spot at various signal to noise ratios. The signal to noise ratio was calculated according to the method shown in figure 8 of this document. The SNR distributions from which these mean SNRs are calculated are shown in Figures 20 and 21 of this document.

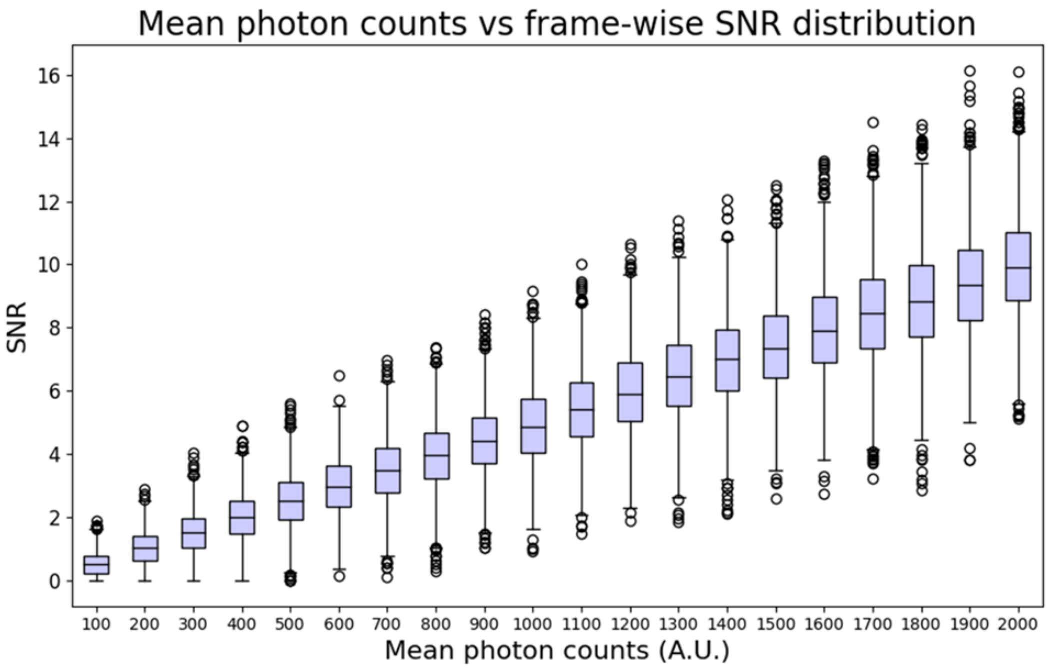


**Figure 20.** Each box plot shows the distribution of SNRs of a single simulated fluorescent spot over 2000 time points. The mean of each box plot is used in figure 19 of this document in which the mislocalization is calculated from the tracked position and known ground truth in each of the 2000 time points.

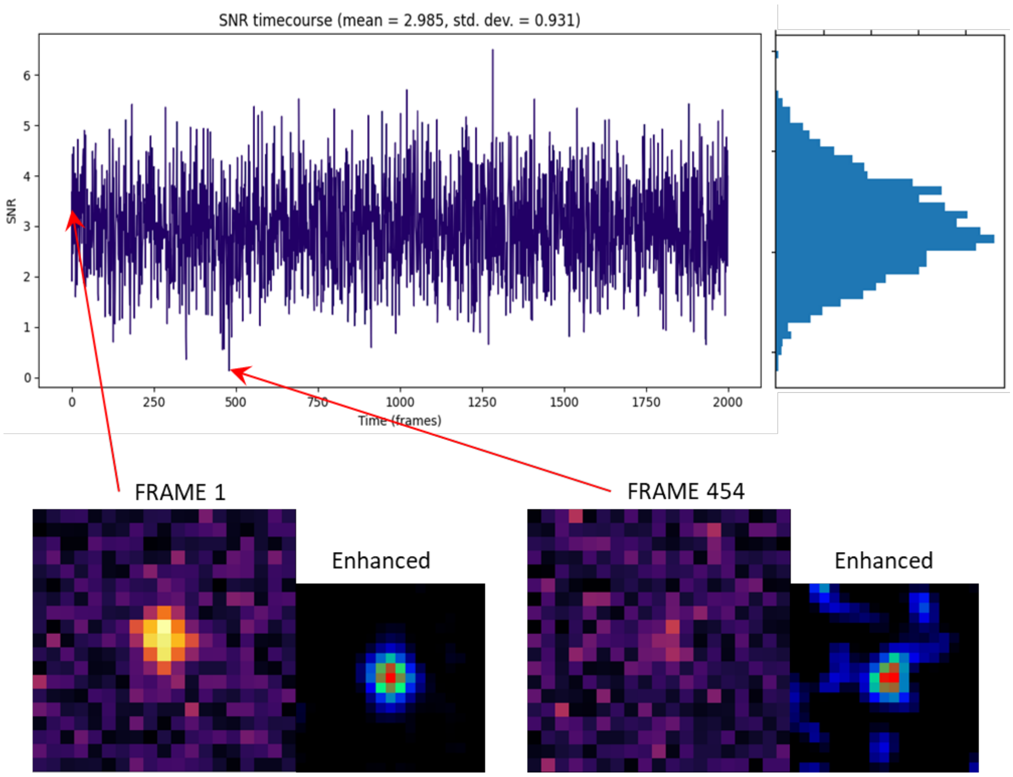


**Figure 21.** A plot of SNR of a spot with a mean simulated photon count of 600 per frame (dee figure 20 above) vs time point over the duration of a 2000 frame synthetic SM movie. Frames of interest are highlighted. Due to the stochastic nature of photon emission some frames (e.g., 454) capture very few photons and result in aberrantly high mislocalization for that frame.

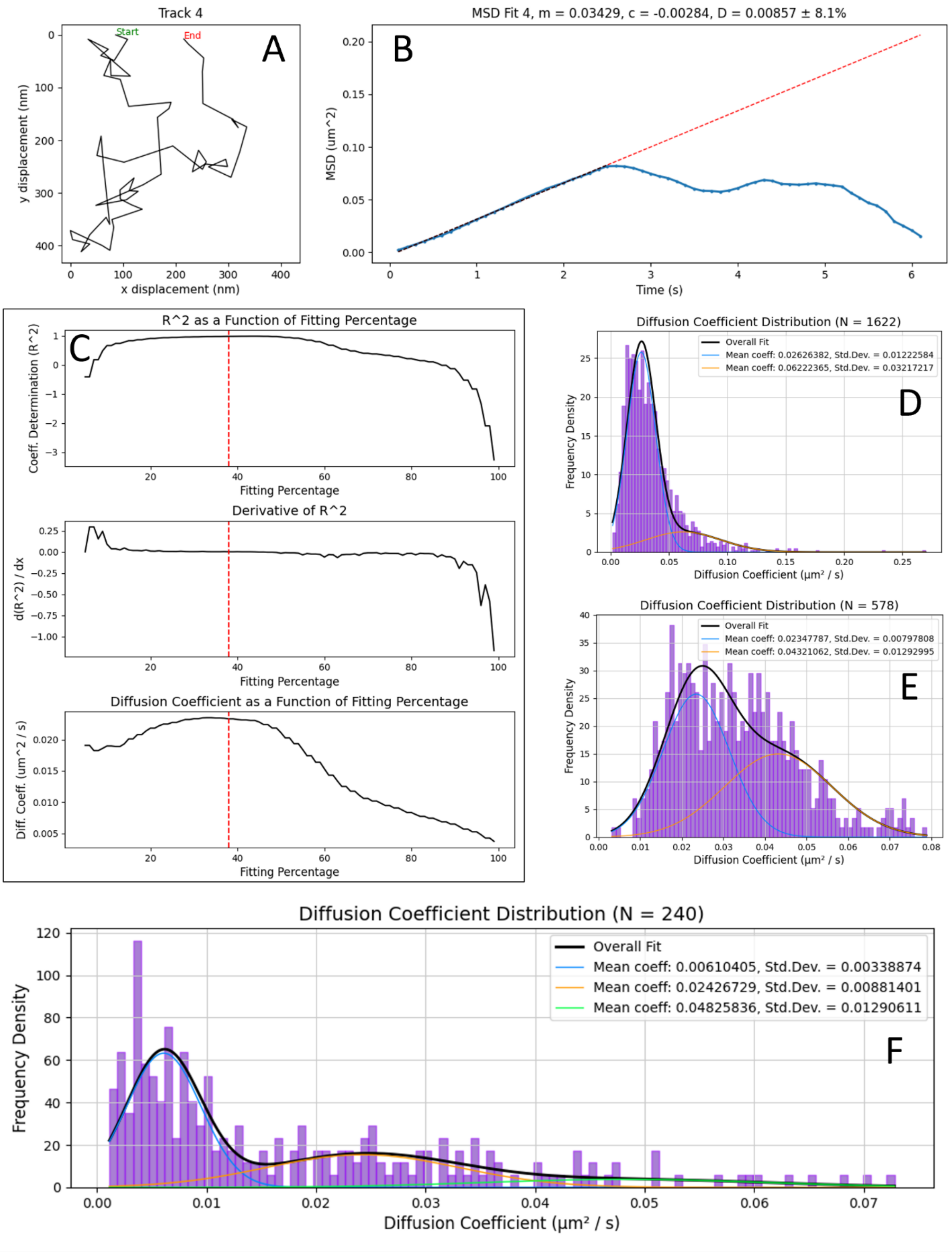

Track 4
Start
End
A
y displacement (nm)
x displacement (nm)
MSD Fit 4, m = 0.03429, c = -0.00284, D = 0.00857 ± 8.1%
B
MSD (um^2)
Time (s)
C
R^2 as a Function of Fitting Percentage
Coeff. Determination (R^2)
Fitting Percentage
Derivative of R^2
d(R^2) / dx
Fitting Percentage
Diffusion Coefficient as a Function of Fitting Percentage
Diff. Coeff. (um^2 / s)
Fitting Percentage
Diffusion Coefficient Distribution (N = 1622)
Overall Fit
Mean coeff: 0.02626382, Std.Dev. = 0.01222584
Mean coeff: 0.06222365, Std.Dev. = 0.03217217
D
Frequency Density
Diffusion Coefficient (μm² / s)
Diffusion Coefficient Distribution (N = 578)
Overall Fit
Mean coeff: 0.02347787, Std.Dev. = 0.00797808
Mean coeff: 0.04321062, Std.Dev. = 0.01292995
E
Frequency Density
Diffusion Coefficient (μm² / s)
Diffusion Coefficient Distribution (N = 240)
Overall Fit
Mean coeff: 0.00610405, Std.Dev. = 0.00338874
Mean coeff: 0.02426729, Std.Dev. = 0.00881401
Mean coeff: 0.04825836, Std.Dev. = 0.01290611
F
Frequency Density
Diffusion Coefficient (μm² / s)

**Figure 22 (Previous page). (A)** An example track from an SM TIRF experiment where a fluorescently labelled single molecule undergoes Brownian motion on a hydrophobic surface. **(B)** The MSD of the track in (A) calculated according to equation 4a. **(C)** A plot of $R^2$ as a function of the proportion of the MSD plot in (B) fitted. $R^2$ in the sci-kit learn implementation can be negative if the correlation between x and y is 0 and the fit is negatively correlated. Below are shown the derivative of $R^2$ as a function of the proportion of fitted points. Below this, the calculated diffusion coefficient as a function of the proportion of fitted points. The red line shows the point at which the MSD begins to deviate from a linear fit. **(D)** Diffusion coefficient histogram of a heterogeneous mixture of simulated single molecules with diffusivities of 0.02 and 0.04 $\mu m^2 s^{-1}$ in equal proportion, fitted with a gaussian mixture model. **(E)** The same data as (D) filtered by estimated error based on the standard deviation in gradients of the fitted proportion of the MSD graph. **(F)** The diffusion coefficient distribution of the experiment described in (A) showing two distinct populations of diffusing molecules. The number of components of the gaussian mixture model has been purposely overfitted to deal with outliers and improve the fit of the main two components.

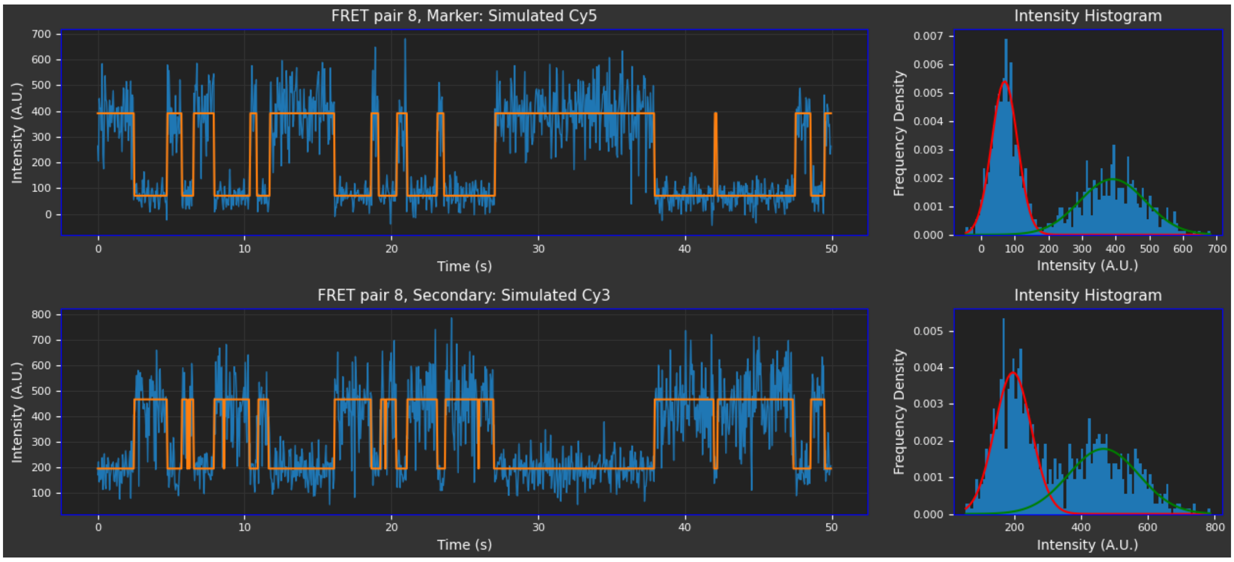


**Figure 23.** Simulated 2-state FRET traces are fitted by finding the mean intensity of each state via a 2-component Gaussian mixture model and the change-points found by scanning a moving average window along the trace and calculating whether the trace switches state at the intersection of the two Gaussian distributions.

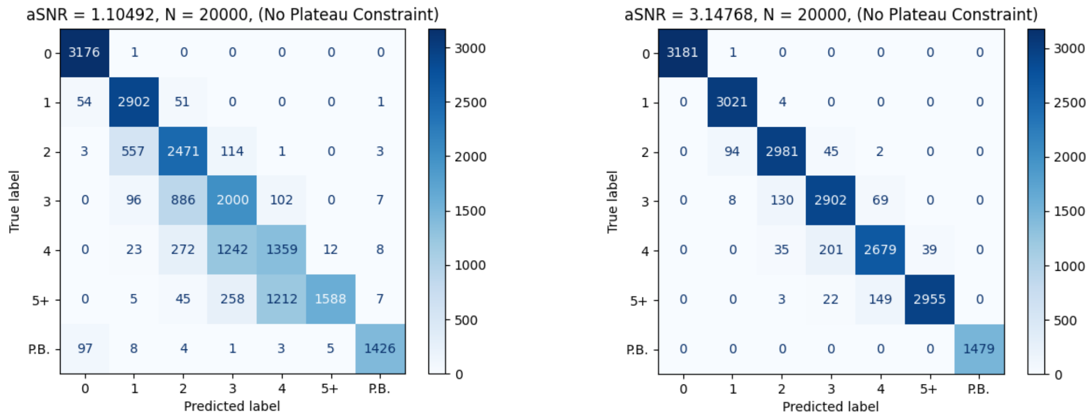


Figure 24. Confusion matrices for accuracy comparisons between simulated traces of mean aSNR 1.1 (left) and 3.1 (right).
